# From raw microalgae to bioplastics: conversion of *Chlorella vulgaris* starch granules into thermoplastic starch

**DOI:** 10.1101/2024.04.17.589749

**Authors:** A. Six, D. Dauvillée, C. Lancelon-Pin, A. Dimitriades-Lemaire, A. Compadre, C. Dubreuil, P. Alvarez, J.-F. Sassi, Y. Li-Beisson, J.-L. Putaux, N. Le Moigne, G. Fleury

## Abstract

Microalgae are emerging as a promising feedstock for bioplastics, with *Chlorella vulgaris* yielding significant amounts of starch. This polysaccharide is convertible into thermoplastic starch (TPS), a biodegradable plastic of industrial relevance. In this study, we developed a pilot-scale protocol for extracting and purifying starch from starch-enriched *Chlorella vulgaris* biomass. From 430.3 ± 0.5 g (dry weight - DW) of microalgae biomass containing 42.2 ± 3.4 % of starch, we successfully extracted 205.8 ± 1.2 g_DW_ of purified starch extract containing 86.9 ± 3.0 % of starch, resulting in a final recovery yield of 98.5%. We have characterized this extracted starch and processed it into TPS using twin-screw extrusion and injection molding. Microalgal starch showed similar properties to those of native plant starch, but with smaller granules. We compared the mechanical properties of microalgal TPS with two controls, namely a commercial TPS and a TPS prepared from commercial potato starch granules. TPS prepared from microalgal starch showed a softer and more ductile behavior compared to the reference materials. This study demonstrates the feasibility of recovering high-purity microalgal starch on a pilot scale with high yields, and highlights the potential of microalgal starch for the production of TPS using industrially relevant processes.

**Graphical abstract:** 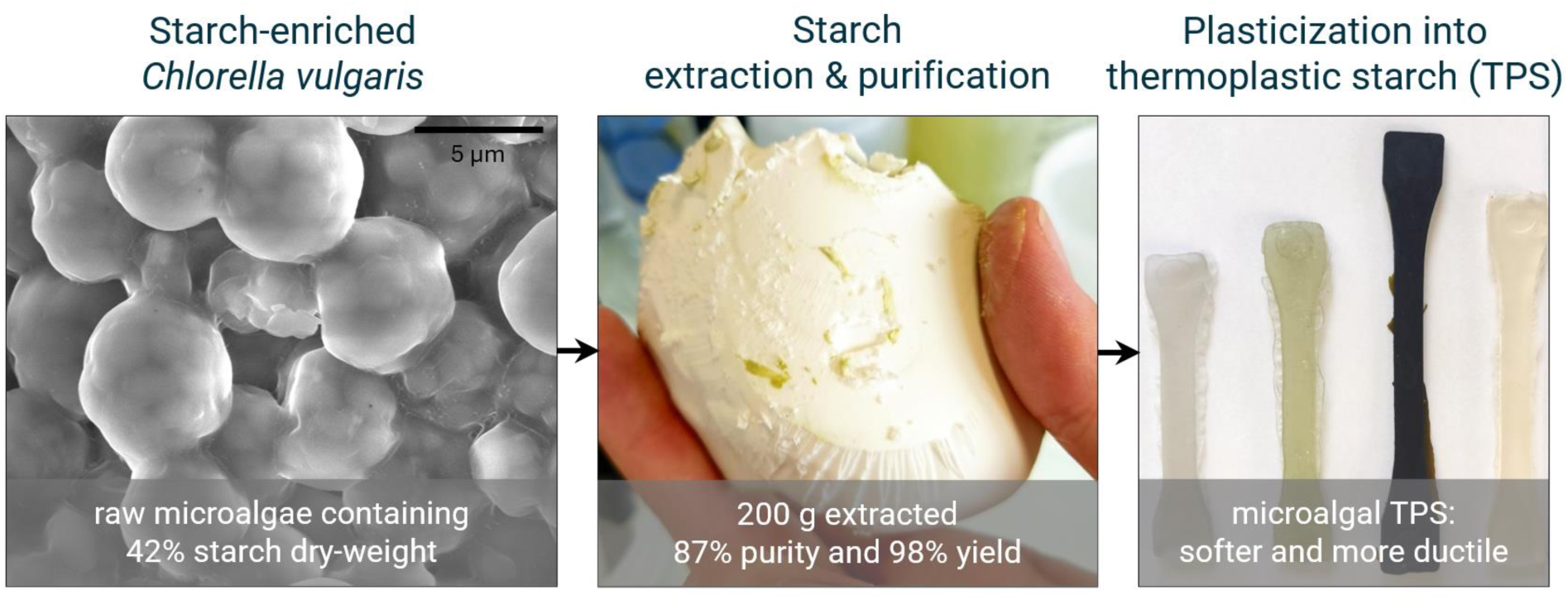

## 1 Introduction

Plastics represent a major source of environmental pollution, with an annual global release of 22 million tons into the environment in 2019 (OECD, 2022). These plastic wastes accumulate in aquatic environments, reaching 139 million tons in 2019, with a staggering projection of 493 million tons by 2060. Plastic debris alter habitats, endanger wildlife, and disrupt the functioning, services, and productivity of ecosystems (Beaumont et al., 2019). Plastic pollution is not only an environmental catastrophe but also a significant economic waste, as ecosystem productivity decreases and billions of dollars of economic value are squandered through single-use and short-lived plastic usage.

The first and most obvious solution to address the plastic pollution problem is to urgently reduce our plastic usage. In parallel, policymakers and industries are pushing towards the use of biobased and biodegradable plastics in different application sectors such as packaging. In this regard, starch emerges as an interesting renewable source for these materials.

Starch is a glucose polysaccharide with α-D-(1→4) and α-D-(1→6) linkages that represents a significant industrial market, with $56B in 2020 (Adewale, Yancheshmeh, & Lam, 2022). Starch can be depolymerized into its glucose subunits to serve as feedstock in fermentation processes to produce biopolyesters, such as poly(lactic acid) (PLA) and polyhydroxyalkanoates (PHAs), or directly plasticized into thermoplastic starch (TPS) with the addition of plasticizers. Yet, the current production of starch from traditional plant crops is unlikely to be sufficient to meet the growing needs of the bioplastics industry, given that starch is mainly intended for food use. This biofeedstock competition with food production could lead to severe biodiversity losses together with alarming socio-economic consequences (Mülhaupt, 2013).

Microalgae could be a solution to increase the global production of starch. Microalgae have a significant biomass production potential. They can grow in wastewater or on non-arable lands, and achieve yields comparable to those of plants (Masojídek, Torzillo, & Koblížek, 2013; Tredici, 2010). In addition, microalgae starch productivity may be similar to plant crops under field conditions (Brányiková et al., 2011). However, one particularity of microalgae compared to plants is the difficulty of accessing the produced starch, as it is enclosed into individual, small, and, depending on the strain, mechanically resistant cells.

Starch extraction from microalgal cells typically consists of three consecutive steps: disruption, separation, and purification. The disruption phase consists in breaking the cell walls to release the intracellular content. For robust microalgal cells, such as *Chlorella vulgaris*, disruption is commonly achieved through chemical hydrolysis, enzymatic treatment, or mechanical stress (Brányiková et al., 2011; Gerken, Donohoe, & Knoshaug, 2013; Yap, Crawford, Dagastine, Scales, Martin, & 2016). Next, the separation phase entails the separation of starch granules from the other components present in the cell lysate (Di Caprio, Amenta, Francolini, Altimari, & Pagnanelli, 2023; Gifuni, Olivieri, Krauss, D’Errico, Pollio, & Marzocchella, 2017; Suarez Ruiz, Baca, van den Broek, van den Berget, Wijffels, & Eppink, 2020). It relies on the high density of starch granules, which allows for an easy recovery by centrifugation. Finally, the starch purification involves the removal of the remaining debris from the starch pellet. In the case of laboratory application, washing is extensive and relies on Percoll gradient centrifugation (Delrue et al., 1992). This degree of purity might not be required for applications such as bioplastics. Other purification approaches have been tested, with acetone or ethanol extraction, and aqueous two-phase systems (Di Caprio et al., 2023; Gifuni et al., 2017; Suarez Ruiz, Baca, et al., 2020; Suarez Ruiz, Kwaijtaal, Peinado, van den Berg, Wijffels, & Eppink, 2020). Yet, these methods are quite complex and fail to achieve both high purity and recovery yield.

Native starch demonstrates thermoplastic properties when processed with plasticizers, elevated temperatures, and shear stress (Zhang, Rempel, & Liu, 2014). Once extracted and purified, the microalgal granular starch can therefore be processed into thermoplastic starch (TPS). Production processes for TPS are well-established and industrialized, notably for packaging films (Polman, Gruter, Parsons, & Tietema, 2021). The main plasticizer used is glycerol, which accounts for 20-30 % of the final weight. However, the residual water naturally present in native starch also acts as plasticizer. Shear and temperature are controlled with a screw extruder, typically in the range of 100-150 °C. The possibility of producing TPS from microalgal starch has already been demonstrated, using the solvent casting technique to produce thin TPS films (Di Caprio et al., 2023). Still, the solvent casting technique is a laboratory-scale method, and the properties of microalgal TPS produced with industrially relevant processes, such as plasticization in twin-screw extruder and injection molding, remains to be evaluated.

The hypothesis of the present study was that starch can be extracted and purified from microalgae in high yield on a pilot scale, and then plasticized into TPS using processes that can be applied on an industrial scale. We adapted and simplified the extraction process of microalgal starch from laboratory to pilot scale, then carried out its extraction, purification, and characterization. Finally, we validated the processability of microalgal starch into TPS by extrusion and injection molding, and evaluated its mechanical properties in comparison with commercial and potato-based TPS.

## 2 Experimental section

### 2.1 Pilot-scale microalgae production for starch extraction

The microalga *Chlorella vulgaris* CCALA924 was cultivated under natural sunlight inside a greenhouse in Saint-Paul-lez-Durance (France). The culture was grown in a 180-L flat panel airlift photobioreactor (PBR) manufactured by Subitec GmbH (Germany). This PBR was operated within a pH range of 7.0-7.5, an air flow of 1500 nL h^-1^ enriched with 2 % of CO_2_ and at a temperature regulated below 30 °C by water aspersion. An adapted Beijrinck medium without NaNO_3_ was used as in (Six et al., 2024). A continuous flow pump of 269 g L^-1^ NaNO_3_ allowed maintaining a concentration of approximately 150 mg L^-1^ NO_3_^-^ during the 6 days of growth phase. The NO_3_^-^ concentration was regularly checked by ion chromatography (940 Professional IC Vario, Metrohm, Switzerland). The pump was stopped on the morning of day 0 to trigger nutrient stress and starch accumulation. The biomass growth in the photobioreactor was monitored both in terms of dry-weight concentration and total carbohydrates (**section 2.2.1**). An 8 % error was considered for dry-weight measurement (Chambonniere et al., 2022). The NO_3_^-^ concentration was null on the afternoon, and the culture was harvested on the morning of day 4. 170 L of culture were concentrated about ten times by SANI membrane filtration (Vibro I, SANI Membranes, Danemark). The pre-concentrated culture was centrifuged (10000 *g*, 10 min) in 1-L buckets (Avanti J-265 XP, Beckman Coulter, USA). The biomass reached a concentration of approximately 40 % (dry weight - DW) and was stored at -20 °C.

### 2.2 Biomass and subfraction characterization

Samples of biomass and their subfractions issued from the extraction process (also referred to as biomass in this section) were frozen at -20 °C and dried (lyophilizer COSMOS 20K, Cryotec, France). The samples were then hermetically sealed and stored in the dark at - 20 °C until analysis. The freeze-dried samples were carefully weighted before analysis with a Mettler Toleto XSR205DU (Greifensee, Switzerland).

#### 2.2.1 Carbohydrate content

The starch accumulation was evaluated by the quantification of total carbohydrates using the protocol developed by Dubois et al. (1956). Briefly, samples of 1–3 mg of lyophilized biomass were digested in 1.25 M H_2_SO_4_ (0.5 mL mg^-1^ DW) at 100.5 °C for 3 h. The digestate was diluted in ultra-pure water to a final volume of 0.5 mL and mixed with 500 µL of 5 % (w/v) phenol solution and 2.5 mL of 95 % H_2_SO_4_. The total carbohydrate content in the suspension was determined by comparing the light absorption at 483 nm with a glucose calibration curve.

#### 2.2.2 Starch content and non-glucose carbohydrates

Starch concentration was determined using the Enzytec Starch kit (R-Biopharm, Germany). Briefly, boiled starches or supernatants were diluted and digested with amyloglucosidase for 15-minutes at 55 °C. The released glucose was calculated though the reduction of NADP^+^ monitored at 340 nm after addition of hexokinase and glucose-6-phosphate dehydrogenase as detailed by the manufacturer. The glucose was converted into starch concentration by applying a factor of 0.9 accounting for the water molecule expulsed during the polymerization of glucose into starch. The concentration of non-glucose carbohydrates was calculated as the difference between glucose and total carbohydrate concentrations. The total carbohydrate concentration was analyzed according to Dubois’ method (see **section 2.2.1**).

#### 2.2.3 Protein content

The protein content in the microalgae was measured using the total nitrogen amount as a proxy, as described by Chambonniere et al. (2022). To evaluate total nitrogen (TN) and total carbon (TC) contents in the biomass, about 10 mg of dry biomass were sampled in the liquid culture, centrifuged, and washed. The resulting pellet was freeze-dried and manually ground with a mortar, to homogenize the particle size. The dry biomass was then suspended in a 50-mL bottle and kept under magnetic stirring to prevent precipitation. The analyses were carried out on the Shimadzu TOC-LCSH system (Shimadzu, Japan) equipped with an 8-channel OCT-L autosampler and a TNM-L Total Nitrogen module. After combustion at 720 °C in a quartz tube, the O_2_ carrier gas (Alphagaz Air 1 provided by Air Liquide, France) directed the sample to infrared detection (NDIR) thermostated at 65 °C, used for total carbon (TC) determination, and to a chemiluminescence detection module for total nitrogen (TN). The flow was set at 150 mLn min^-1^. The protein fraction in the algal biomass was computed assuming a nitrogen-to-protein ratio of 5.04, as reported for *Chlorella vulgaris* (Templeton & Laurens, 2015).

#### 2.2.4 Lipid content

The lipid content and profile were assessed by total fatty acid analysis using GC-FID (GC-2010 Pro AOC-20i/AOC-20s, Shimadzu™, Japan). Lyophilized biomass samples (2–10 mg) underwent transmethylation to produce fatty acid methyl esters (FAMEs). Briefly, the samples were suspended in 3 mL of a transesterification agent (1.25 M hydrogen chloride in methanol, Ref: 17935, Supelco, USA) and combined with 0.2 mL of a 3 mg L^-1^ TAG C15:0 solution (Tripentadecanoin, reference T4257, Sigma-Aldrich™, USA) in anhydrous hexane 95 % (296090, Sigma-Aldrich™, USA) as an internal standard. This mixture was incubated at 85 °C for 1 h. Subsequently, 3 mL of HPLC-grade hexane was added, followed by vigorous mixing with 1 mL ultra-pure water. After centrifugation (524 *g*, 5 min), the upper hexane phase was collected for GC analysis. The total lipid content was calculated by comparing the sum of peak areas to that of the internal standard. Compound identification utilized retention times and validation with a set of standards of 37 FAMEs (ref. CRM47885, Supelco, USA).

#### 2.2.5 Pigment content

The pigment content was measured by the colorimetric method of Wellburn (1994) after pigment extraction in dimethyl sulfoxide (DMSO). Briefly, pure DMSO was added in samples of 0.5–1.5 mg of freeze-dried biomass to reach 2 mL mg^-1^ DW. Samples were heated in the dark at 60 °C for 40 min. After centrifugation, the optical density of supernatant was measured at 450, 649 and 665 nm in 1-cm lightpath cuvette (ref. 11602609, ThermoFisher, USA) using an UV-Vis Epoch2 (BioTek Instruments, USA). The concentrations in chlorophyll *a*, chlorophyll *b* and total carotenoids were calculated according to the equations reported by Wellburn (Wellburn, 1994):

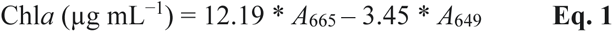

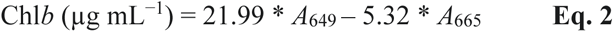

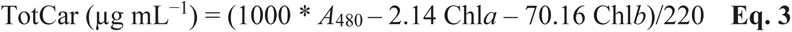

with Chl*a* and Chl*b* the chlorophyll *a* and *b* concentrations, respectively, TotCar the total carotenoid concentration, and A_665_, A_649_, and A_480_ the absorbances at 665, 649, and 480 nm, respectively. The total chlorophyll content was calculated as the sum of chlorophyll *a* and chlorophyll *b* contents.

#### 2.2.6 Ash content

At least 100 mg of pre-weighted samples were placed in a muffle oven (Model LE, Nabertherm, Germany). The temperature was gradually increased to 550 °C for 1 h and was then maintained for 2 h. Ashes from the pre-weighted samples were subsequently recovered and weighted.

### 2.3 Assessment of cell rupture

Samples of biomass were stored at -20 °C without freeze-drying to preserve the integrity of cell structures. Thawed biomass samples were diluted to the desired concentration in pure water and cells were disrupted by high-pressure homogenization (HPH, CF2 Cell Disruptor, Constant Systems Ltd., England) at 250 MPa. The biomass broth was retrieved and submitted to HPH for several repeated cycles. The cell concentration was measured in triplicate with a particle counter (Multisizer 4 Counter, Beckman Coulter, USA) after each disruption cycle. Particles outside the typical cell size range of *Chlorella vulgaris*, from 2.6 to 10 µm, were excluded. The cell rupture was calculated as the ratio of the disappeared cell concentration (difference in present and initial cell concentration) to the initial cell concentration.

### 2.4 Starch characterization

#### 2.4.1 Molecular structure

Starch granules were extracted from the microalgae after complete cell rupture by HPH at 250 MPa for 5 cycles. The biomass broth was centrifuged for 5 min at 4500 *g* to pellet the starch granules. The supernatant was discarded and the pellet was resuspended in 1 mL of Percoll^®^ (GE Healthcare Bio-Science AB, Sweden). After strong agitation followed by centrifugation, the supernatant of biomass debris and Percoll^®^ was carefully removed. The starch pellet was rinsed in pure water and freeze-dried. Amylose and amylopectin were separated by gel permeation chromatography on a Sepharose CL-2B column (0.5 cm inner diameter, 65 cm length) eluted in 10 mM NaOH at aflow rate of 12 mL h^-1^ and the wavelength at the maximum absorbance of the iodine polysaccharide complex (λ_max_) was measured for each 300-μL fraction as described by Delrue et al. (1992).

#### 2.4.2 Scanning electron microscopy (SEM)

A droplet of a dilute starch granule suspension was air-dried on copper tape fixed on a metallic stub. After a coating of the surface with Au/Pd in a Safematic CCU-010-HV sputter coater, secondary electron images were recorded with a Thermo Scientific FEI Quanta 250 microscope equipped with a field-emission gun and operating at 2.5 kV. Injection-molded specimens were cryofractured in liquid nitrogen, their cross-section surface was coated with carbon (Carbon Evaporator Device CED030, Balzers), and observations were performed with a Thermo Scientific FEI Quanta 200 FEG microscope operating at 12.5 kV.

#### 2.4.3 Small- and wide-angle X-ray scattering (SAXS and WAXS)

An aqueous suspension of native starch granules was centrifuged. The pellet was allowed to desorb in a chamber maintaining a 95 % relative humidity (r.h.) for 5 days and poured into a 1-mm outer diameter glass capillary that was flame-sealed and X-rayed in transmission in vacuum (Ni-filtered CuKα radiation, λ = 0.1542 nm). WAXS two-dimensional patterns were recorded on Fujifilm imaging plates at sample-to-detector distances of 4.5 cm. For the SAXS analysis, a capillary containing starch granules in excess water was X-rayed at a distance of 29 cm. The exposed imaging plates were read offline with a Fujifilm BAS 1800-II bioanalyzer. Scattering profiles were calculated by radially averaging the 2D patterns.

#### 2.4.4 Differential scanning calorimetry (DSC)

Starch granules (*ca.* 10 mg) were poured into a 60-mL stainless steel pan to which 30 mg of deionized water was added. The pan was hermetically sealed, accurately weighted, and allowed to equilibrate overnight. The sample pan was scanned against a pan containing 40 mg water in a TA Instruments Q200 differential scanning calorimeter. The sample was heated from 20 to 140 °C, then cooled down to 20 °C, and immediately reheated to 140 °C, all steps carried out at a rate of 5 °C.min^-1^.

#### 2.4.5 Granulometry

The particle size distribution was evaluated by light scattering using a Horiba LA950 laser granulometer (0.05 - 3000 µm size range). The starch granule suspension was added dropwise in the circulation loop to achieve an obscuration rate between 10 and 70 %. The size distribution was determined using Mie’s theory, assuming a refractive index of 1.55 for starch and 1.33 for water, and calculating a particle diameter equivalent to that of a sphere of similar volume.

#### 2.4.6 Statistical analysis

Statistical significance was assessed using the ANOVA test with Prism 9 (Graphpad Software, LLC). When the null hypothesis was rejected (p < 0.05), data were further analyzed using Tukey’s multiple comparison test. Unless otherwise indicated , the following results are presented as the average of the technical replicate (n = 3), while the error bars account for the standard deviation.

### 2.5 Starch extraction, plasticization and characterization

#### 2.5.1 Extraction from microalgal biomass

The thawed starch-enriched biomass was diluted to 10 % DW in pure water. The suspended cells were disrupted by HPH at 250 MPa for 5 cycles. The cell lysate was centrifuged at about 16000 *g* for 10 min at 5 °C (Avanti J-265 XP, Beckman Coulter, USA). The starch pellet was recovered, resuspended at a concentration of 5 L of pure water per kg of wet starch with magnetic stirring, and centrifuged at 15900 *g* for 5 min at 5 °C. This rinsing was repeated twice. The purified starch was freeze-dried for conservation purpose.

#### 2.5.2 Starch materials and control TPS

Three types of materials, *i.e.* potato starch, microalgal starch, and raw biomass, were used to conduct the plasticization study. Potato starch was a commercial grade from Avebe, Netherlands, of which some characterizations can be found in literature (Avebe, 2017; Kim, Wiesenborn, Lorenzen, & Berglund, 1996; van Soest et al., Hulleman, de Wit, & Vliegenthart, 1996; Zhang et al., 2014). Microalgal starch refers to the purified starch from the pilot-scale process, after freeze-drying (**section 2.5.1**). Due to a higher dryness, microalgal starch was also complemented with 10 % water (final weight). Raw biomass refers to the entire starch-enriched microalgae cells that were used for microalgal starch extraction, in their freeze-dried form. The water content in the three materials was analyzed with an infrared moisture analyzer MA35 (Sartorius, Germany), using approximately 1 g per measurement and three replicates per material, after three days of moisture content equilibration in a chamber at 50 % r.h. Pure glycerol was added to the three materials to achieve a starch material / glycerol ratio of 70 % / 30 %, and the mixture was kept at rest overnight at 4 °C. Finally, commercial TPS (NP WS 001, NaturePlast, France) was used as a control for studying TPS processing and properties.

#### 2.5.3 Optical microscopy observations of starch plasticization

The plasticization of the three materials (potato starch, microalgal starch and raw biomass) in glycerol under static conditions (*i.e.* without shearing) was observed with an optical microscope in transmitted light (Laborlux 11 POL S, Leitz, Germany) equipped with a heating stage (LTS420, Linkam, UK). The use of polarized light and gypsum retardation plate enabled the observation of birefringent structures. Videos were recorded with a digital camera (Leica DFC 420, 5 megapixel CCD) piloted by the Replay software (Microvision Instruments, France). Samples of potato starch, microalgal starch and raw biomass were placed under two glass plates in the presence of a large excess of glycerol. A heating cycle at 5 °C min^-1^ was applied to the mixture up to 160 °C.

#### 2.5.4 Extrusion and injection of TPS

The starch material and glycerol mixtures were split in 12-14 g batches, sufficient to fill the mixing chamber of the twin-screw extruder (microcompounder model MC5, Xplore, Netherlands). Twin-screw extrusion (TSE) is a well-established and upscalable method for processing thermoplastics, used to mix, additivate and/or plasticize polymers, creating homogeneous blends. The starch material mixed with glycerol was directly plasticized at temperatures of 120 and 140 °C, with a screw speed of 100 rpm and mixing time of 2 min. Commercial TPS pellets and plasticized starch extrudates collected in each processing condition were injected into tensile specimens with a sequential injection molding machine (Zamak Mercator, Poland) in accordance with ISO 527-2 1BA specifications. The sheath temperature was 140 °C, the mold temperature 40-45 °C and the injection pressure 5.5 bar.

#### 2.5.5 Tensile tests

Tensile properties of the resulting TPS specimens were measured according to the ISO 527 standard with a Zwick TH 010 testing machine. Before testing, the injected specimens rested for three days in a room regulated at 23 °C and 45 % r.h., and tensile tests were performed with a temperature and humidity varying between 23-24 °C and 41-46 % r.h., respectively. For each sample, the thickness and width were precisely measured on three different regions of the specimen, and the mean values were used for stress calculations. The parameters used to establish the stress-strain curves and determine Young’s modulus, ultimate tensile strength and strain at break, were a pre-stress of 0.5 N and a strain rate of 60 % gauge length per min. At least 7 samples per blend were tested.

## 3 Results and Discussion

### 3.1 Extraction of the starch-enriched microalgae

The first objective consisted in obtaining an important quantity of microalgae biomass enriched in starch. To do so, a culture of *Chlorella vulgaris* CCALA924 was grown under natural sunlight in a 180-L flat panel PBR, and enriched in starch by triggering nitrogen starvation, as described in the Experimental section. The evolution of biomass and total carbohydrate concentration is reported in **Figure 1**. At the end of the culture, a biomass containing 42.2 ± 3.4 % DW of starch was retrieved.

**Figure 1:**
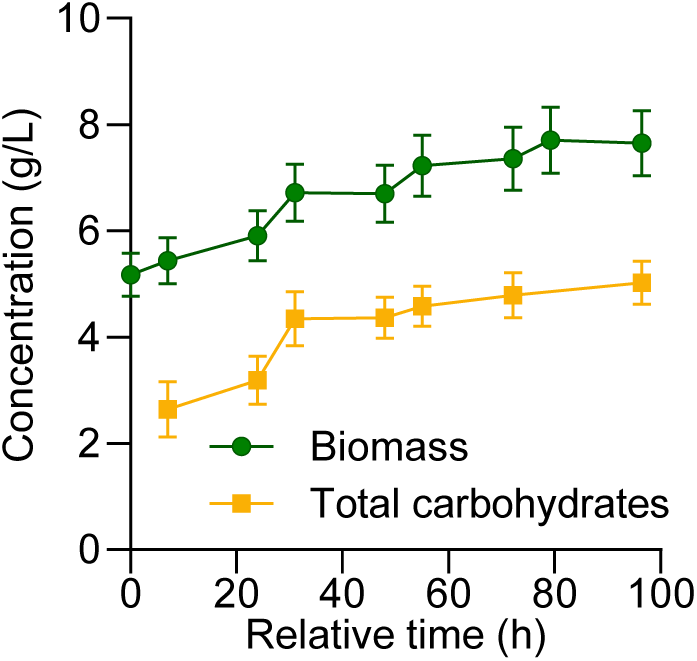
Growth of the microalgae biomass after induction of starch accumulation by nitrogen deprivation. Technical replicates: biomass n=1, carbohydrates n=3. For biomass concentration, an 8 % error was considered, as described in the Experimental section.

#### 3.1.1 Adaptation of the extraction protocol

Current starch extraction methods from microalgae are typically conducted at laboratory scale. The different protocols either exhibit purity or extraction yield below industrial standards (Di Caprio et al., 2023; Gifuni, 2017), involve complex process (Suarez Ruiz, Baca, et al., 2020), or require expensive products (Delrue et al., 1992). One objective of our work was to adapt the expensive yet simple, efficient protocol described by Delrue et al. (1992) and developed in (Vonlanthen, Dauvillée, & Purton, 2015) to the pilot scale by changing the cell disruption method from French press to semi-continuous high pressure homogenization (HPH) and by replacing the starch purification with the expensive Percoll by only repeated water rinsing. The comparison of cell disruption efficiency with French press and HPH is presented in **Figure S1**, and HPH disruption efficiency is shown in **Figure 2** for different disruption cycles and biomass concentrations. The pictures showing the effect of water rinsing on starch purification are shown in **Figure S2**.

**Figure 2:**
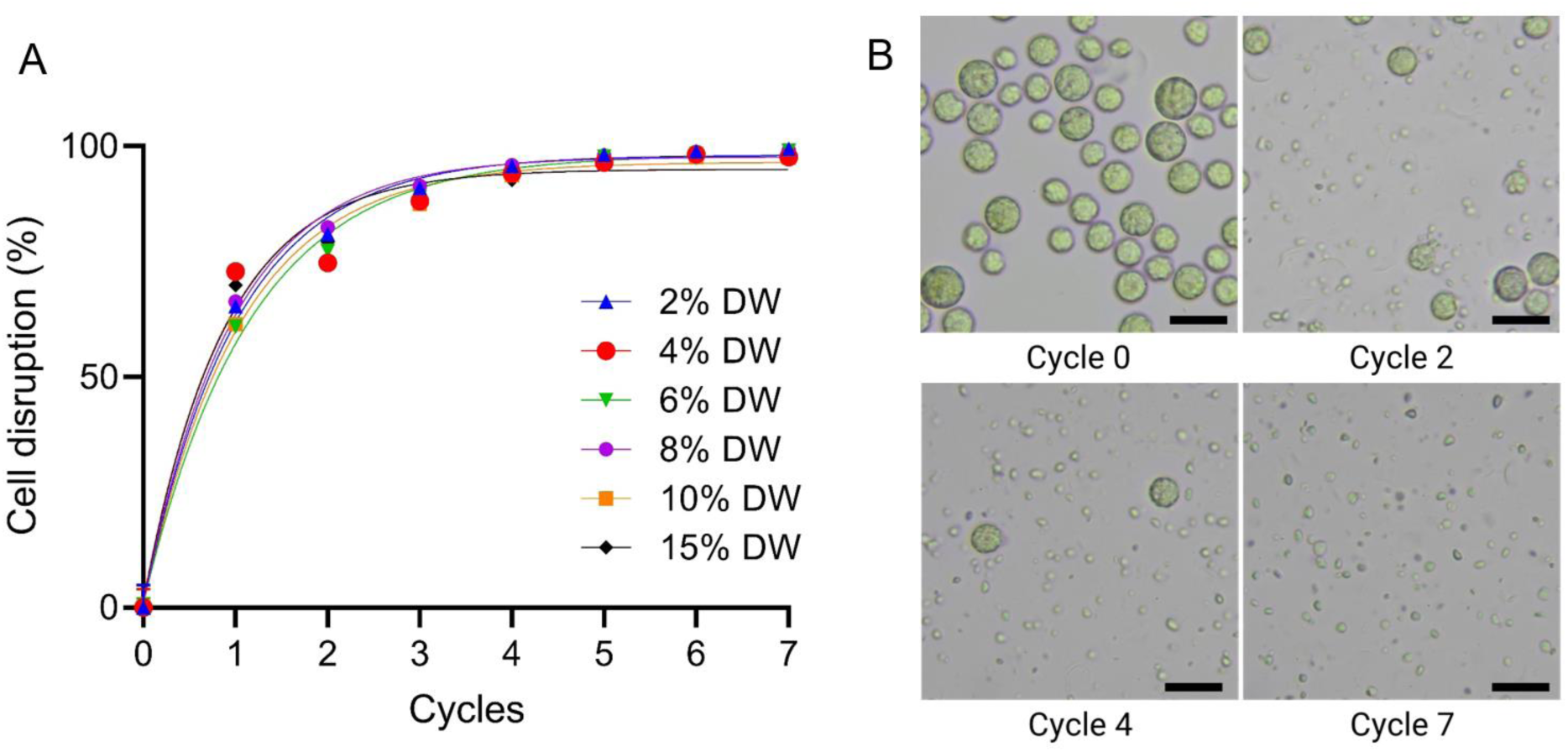
Disruption of microalgal cells during the pressure cycles: (A) efficiency of high-pressure homogenization at 250 MPa for different biomass concentrations; (B) Disruption of the microalgal cells after the HPH cycles on a 4 % DW biomass. Scale bars correspond to 10 µm. The cell disruption was analyzed in technical triplicates.

Cell disruption occurred similarly with French press and HPH at the same pressure (**Figure S1**). The biomass concentration was attested to have no effect on the cell disruption efficiency with HPH, up to a concentration of 15 % DW (**Figure 2**). This result was also demonstrated by Yap et al. which argued for a maximal concentration of 25 % DW for another strain of *Chlorella vulgaris* (Yap et al., 2015). The release of starch granules and cell wall fragments in the suspension was confirmed by optical microscopy (**Figure 2**). More than 99 % of the cells were disrupted after 5 cycles of HPH at 250 MPa. In addition, replacing the Percoll addition with three successive water rinsing resulted in a visible removal of cell debris from the starch fraction (**Figure S2**).

#### 3.1.2 Pilot-scale starch extraction

After the adaptation of the starch extraction protocol, we aimed at recovering at least 200 g of purified microalgal starch to perform plasticization tests. The starch was extracted from the microalgae cells following the starch extraction protocol presented in **Figure S3** and **Figure S4**. First, the microalgae cells were diluted to 10 % DW in water and then disrupted by HPH at 250 MPa for 5 cycles. Following the disruption, the centrifugation of the cell lysate resulted in the separation of three fractions: a supernatant and a pellet divided into two parts. The upper part of the pellet was green and doughy, and the bottom part was white and dense. At this stage, the dry weight content of supernatant, green pellet, and starch pellet were of 4.2, 13.1 and 55.8 % of wet weight, respectively. The composition of these three fractions is reported in **Table 1**. The starch fraction was then manually recovered from the other two fractions, and subsequently purified by three cycles of water addition, starch resuspension, centrifugation, and supernatant removal. The rinsing supernatants decreased in turbidity and coloration after each rinsing cycle, until no chlorophyll absorption peak was detected in the last rinsing supernatant (**Figure 3**). The purification by water rinsing reduced the non-glucose carbohydrates and lipid content, while the starch content slightly increased (**Table 1**).

**Figure 3:**
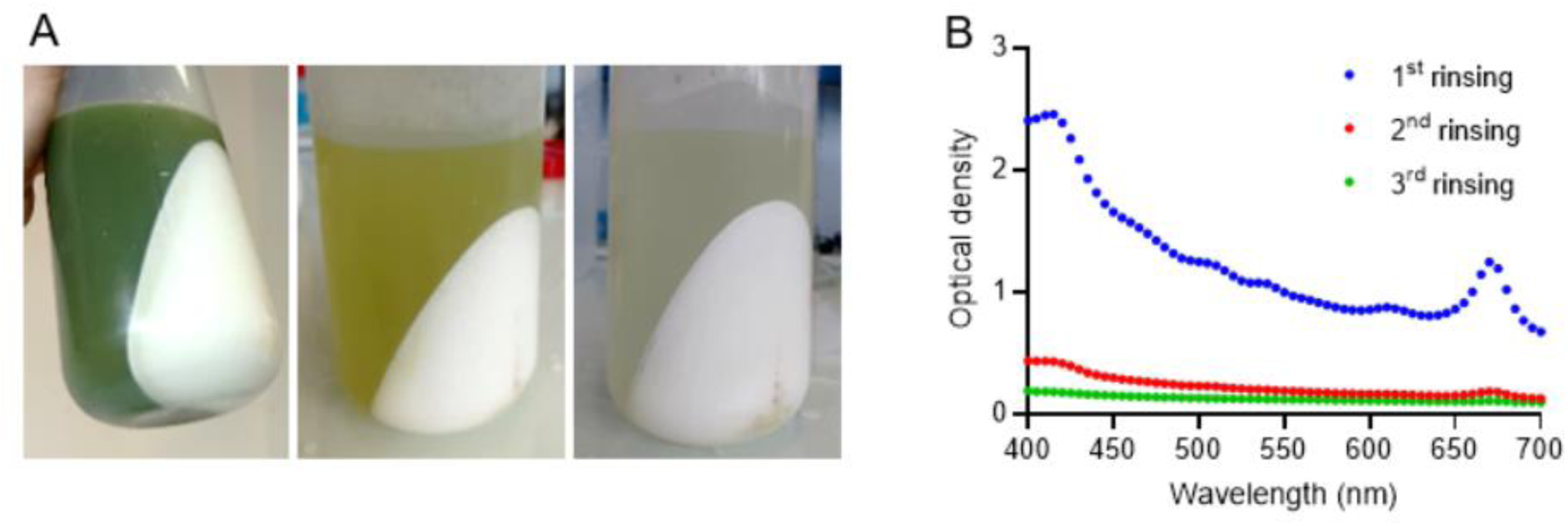
Starch purification with successive water rinsing. (A) Pictures of pellet and supernatant after each rinsing step. (B) Absorbance profile of supernatants after each water rinsing step.

**Table 1:**
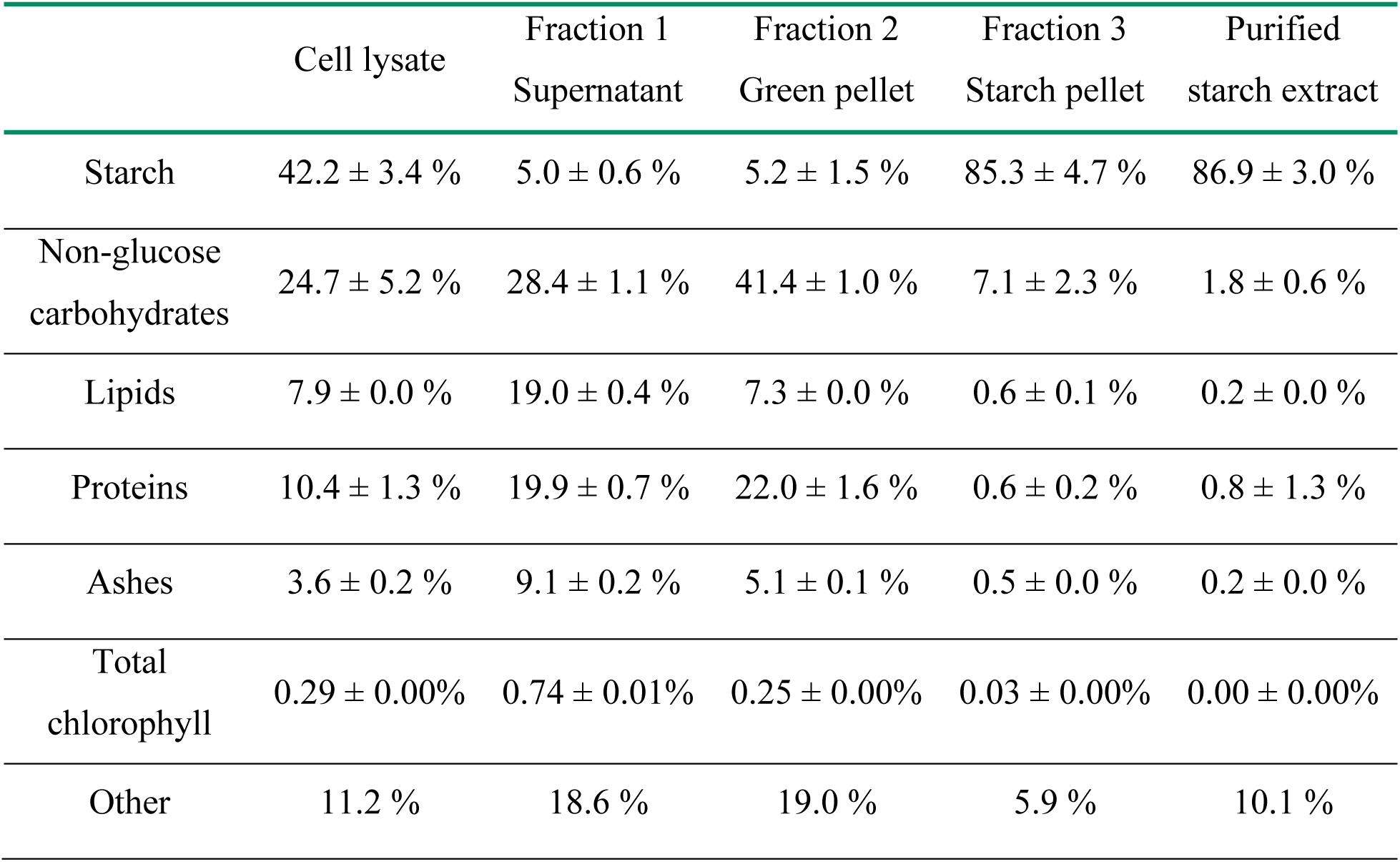
Content in class of compounds for initial cell lysate, separated fractions, and purified starch. The composition of non-disrupted biomass was very similar to the cell lysate, except for starch and non-glucose carbohydrates, that could not be determined in entire cells.

|  | Cell lysate | Fraction 1<br>Supernatant | Fraction 2<br>Green pellet | Fraction 3<br>Starch pellet | Purified<br>starch extract |
| --- | --- | --- | --- | --- | --- |
| Starch | 42.2 ± 3.4 % | 5.0 ± 0.6 % | 5.2 ± 1.5 % | 85.3 ± 4.7 % | 86.9 ± 3.0 % |
| Non-glucose<br>carbohydrates | 24.7 ± 5.2 % | 28.4 ± 1.1 % | 41.4 ± 1.0 % | 7.1 ± 2.3 % | 1.8 ± 0.6 % |
| Lipids | 7.9 ± 0.0 % | 19.0 ± 0.4 % | 7.3 ± 0.0 % | 0.6 ± 0.1 % | 0.2 ± 0.0 % |
| Proteins | 10.4 ± 1.3 % | 19.9 ± 0.7 % | 22.0 ± 1.6 % | 0.6 ± 0.2 % | 0.8 ± 1.3 % |
| Ashes | 3.6 ± 0.2 % | 9.1 ± 0.2 % | 5.1 ± 0.1 % | 0.5 ± 0.0 % | 0.2 ± 0.0 % |
| Total<br>chlorophyll | 0.29 ± 0.00% | 0.74 ± 0.01% | 0.25 ± 0.00% | 0.03 ± 0.00% | 0.00 ± 0.00% |
| Other | 11.2 % | 18.6 % | 19.0 % | 5.9 % | 10.1 % |

The final composition of the purified starch was close to the typical composition reported for granular starch (2 % lipids, 0.6 % proteins, and 0.4 % minerals) ( Zhang et al., 2014). Finally, from 430.3 ± 0.5 g_DW_ of microalgal biomass containing 42.2 ± 3.4 % of starch, 205.8 ± 1.2 g_DW_ of purified starch extract containing 86.9 ± 3.0 % of starch were extracted. At the end of the process, we obtained 178.8 g of starch from the 181.5 g of starch sheathed in the microalgae cells, for a final recovery yield of 98.5 %. A second starch extraction was conducted on the microalgal biomass issued from the same culture, resulting in a similar starch recovery yield (data not shown), with a final extraction of approximately 80 g pure starch.

This starch extraction process not only offered an efficient starch recovery, but it also opened interesting biorefinery perspectives. Indeed, valuable microalgal compounds such as proteins and lipids were also extracted during the separation phase. The supernatant was rich in lipids and proteins (19.0 ± 0.4 %_DW_ lipids, 19.9 ± 0.7 %_DW_ proteins), while the green pellet contained a similar protein content but less lipids (7.3 ± 0.0 %_DW_ lipids, 22.0 ± 1.6 %_DW_ proteins) (**Table 1**). Interestingly, the supernatant was the fraction containing most of the lipids and proteins initially present in the cell lysate, with 79.5 ± 1.5 % and 61.1 ± 7.9 % recovery, respectively (**Table S1**). Moreover, in a previous study, the lipid profile of *Chlorella vulgaris* CCALA924 was shown to contain a majority of polyunsaturated fatty acids and only around 20 % of saturated fatty acids (Six et al., 2024), making the lipid fraction attractive for feed or food applications. However, the supernatants and green pellets were ultimately mainly composed of water containing a dry mass of only 4.2 and 13.1 %, respectively. Moreover, the recoverability of lipids and proteins from these fractions containing a large amount of micronized cell debris is still an open question. In addition to creating micronized debris (Harrison, 1991), the multiple passes in HPH might be accompanied by the emulsion of lipids which may further hinder their downstream processing.

However, despite the efficiency of the starch extraction process, the techniques employed are highly energy-demanding, especially for the cell disruption, and could be prohibitive for industrial applications. Indeed, Yap et al. evaluated that if a *Chlorella vulgaris* culture at 10 % DW and containing 30 % of lipids was disrupted with one pass of HPH at 150 MPa, the energy consumption of HPH would be equivalent to 37 % of the energy effectively accessible in the extracted lipids (Yap et al., 2015). In this work, we concluded on the necessity to run 5 cycles of HPH at 250 MPa to extract the totality of the starch. Therefore, the HPH disruption appeared as highly inefficient regarding the required energy. Other disruption techniques, such as enzymatic cell wall disruption, could reduce the energy footprint of the extraction process.

On the other hand, the water footprint of the process was more advantageous. Here, the three rinsing steps used approximately 13 L of water per kg of dry biomass in total, equivalent to 26 L of water per kg of dry-pure-starch. Tran et al. have shown that cassava starch extraction required from 9.8 to 20.8 m^3^ of water per ton of starch depending on the presence of water recycling in the process (Tran et al., 2015). Therefore, the water footprint of our microalgal starch extraction at pilot scale looked similar to the water footprint of industrial-scale extraction of plant starch.

### 3.2 Morphology and structure of microalgal starch

SEM observations of the purified microalgal starch indicated that the granule integrity was preserved throughout the extraction process (**Figure S5**). The starch granules exhibited a mean diameter of 1.5 µm, as determined by laser granulometry (**Figure 4A**). Although this size was relatively small compared to plant starch, it is typical of green microalgae (Di Caprio et al., 2023). Interestingly, the microalgal starch contained 13.0 % amylose (**Figure 4B**), a relatively low content for storage-type starch usually obtained under nitrogen-deprived conditions (Findinier et al., 2019).

**Figure 4:**
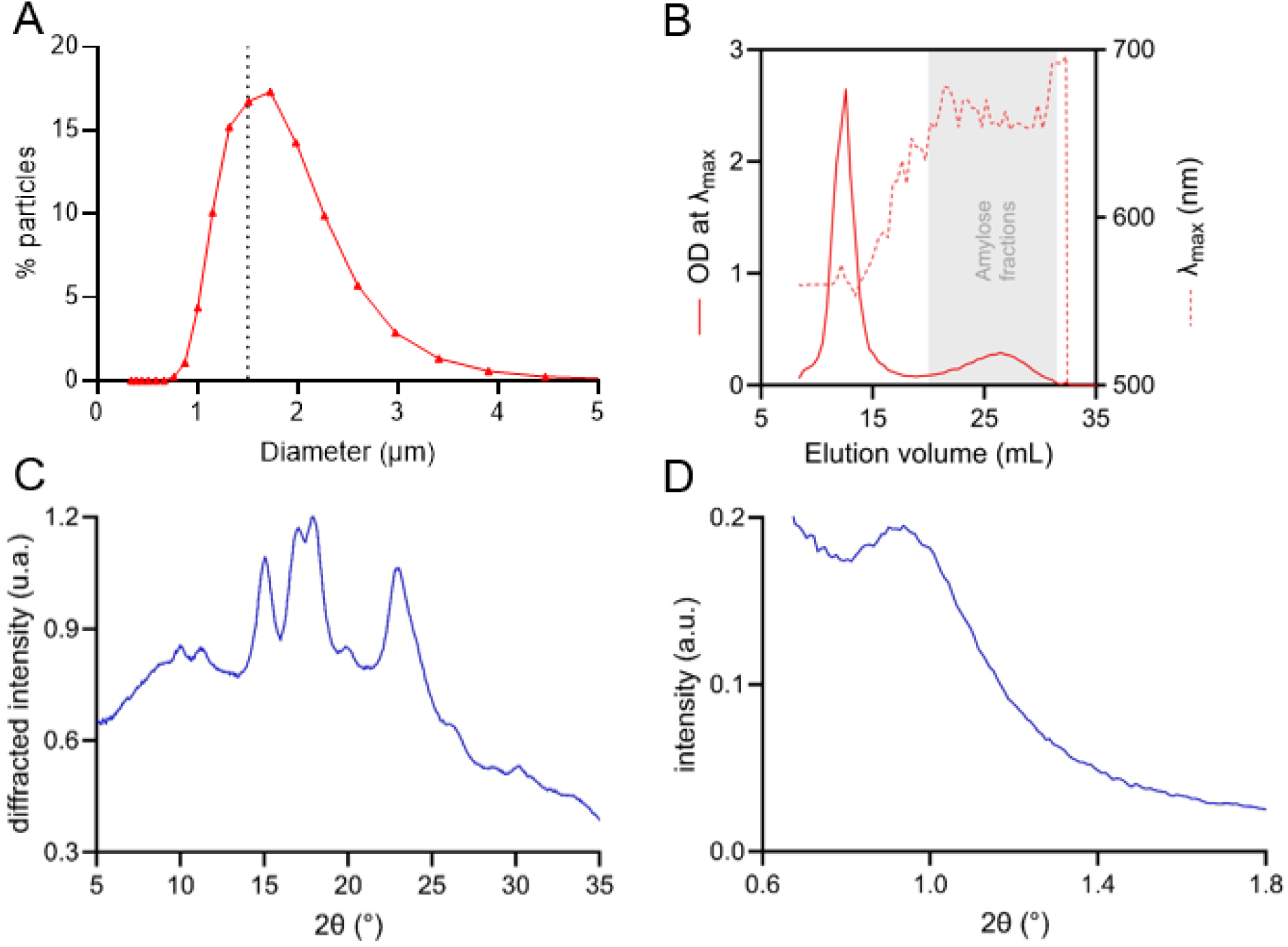
Structural characterization of the extracted microalgal starch. (A) Size repartition of starch granules analyzed with laser granulometry. (B) Separation of amylopectin and amylose on CL-2B chromatography. The maximum absorbance of the polysaccharide/iodine complex (full line) was measured between 500 and 700 nm for each fraction and the wavelength at maximal absorbance (λ_max_ in nm, dotted line) was determined. The gel permeation chromatography excluded the high molecular weight amylopectin (1^st^ peak; between 8 and 15 mL of elution); while lower molecular weight amylose was eluted later (2^nd^ peak, after 18 mL of elution, grey area). (C) WAXS profile of hydrated microalgal starch granules, and (D) SAXS profile of granules in excess water.

The WAXS profile of the microalgal starch revealed an A-type crystal structure, with peaks located at 2θ = 9.9, 11.2, 15.1, 17.0, 17.9, 20.0, and 22.9° (**Figure 4C**). The peak in the SAXS profile, at 2θ = 0.93°, indicated an interlamellar repeat of 9.5 nm for amylopectin (**Figure 4D**). Overall, the characteristics of the extracted starch fall within the typical range observed for Chlorella starch (Gifuni et al., 2017).

The DSC thermogram of microalgal starch during heating in excess water showed an endothermic peak at 65 °C corresponding to the gelatinization of the granules (**Figure 5)**. This temperature falls within the range of expected values, as it was shown that microalgal starch gelatinization temperature varied between 65 and 75°C (Izumo et al., 2007). During cooling, a small exothermic peak at 75 °C was detected that was attributed to the crystallization of amylose with residual lipids. On second heating, a broad and faint endothermic peak appeared from 100 °C, which could correspond to the melting of the amylose-lipid complexes. The DSC thermogram of maize starch, which also displays A-type crystallinity and contains lipids, usually displays the same trend (Arik Kibar, Gönenç, & Us, 2014; Eliasson, Finstad, & Ljunger, 1988). The DSC of microalgal starch did not differ much compared to potato starch which does not contain lipids (**Figure 5**). Overall, the thermal behavior of microalgal starch granules in excess water was similar to that of standard starch.

**Figure 5:**
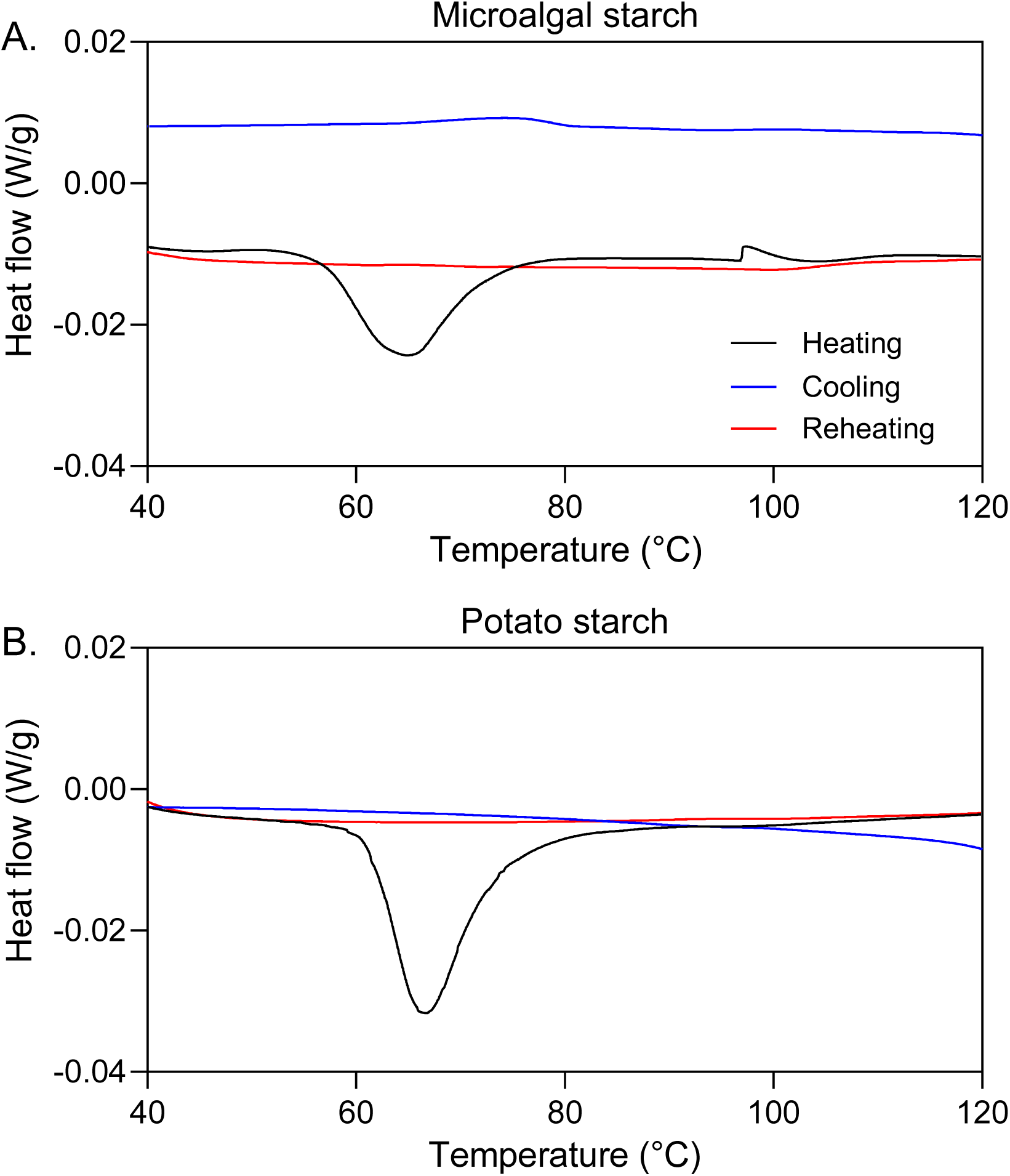
Differential scanning calorimetry thermograms of microalgal and potato starches. The changes of physical properties along the temperature are revealed by the changes of heat flow in microalgal starch (A), and potato starch (B). The unexpected peak appearing around 100 °C for microalgal starch is probably due to the occurrence of a bubble.

### 3.3 Plasticization of microalgal starch and characterization

The plasticization of the purified microalgal starch into TPS was compared with that of two control starches. The first one was a commercial grade of potato starch, widely employed in industrial thermoplastic-starch production. In contrast, the second control consisted of our freeze-dried raw microalgal biomass, containing intact starch-enriched cells. Although it has been shown that the use of entire cells resulted in incomplete plasticization (Mathiot, Ponge, Gallard, Sassi, Delrue, & Le Moigne, 2019), the comparison with purified microalgal starch should provide insights into the relevance of the extraction and purification process with regard to plasticization. Furthermore, the direct plasticization of entire cells might be an interesting alternative route to limit processing steps in the production of microalgae-based TPS.

#### 3.3.1 Plasticization monitored by temperature-controlled optical microscopy

Initially, the plasticization capacity of the three materials was investigated in the presence of excess glycerol and without the application of shear stress. A small quantity of dry material was introduced between two glass plates into a large volume of glycerol and gradually heated to 160 °C. The behavior of starch granules under heating was observed *in situ* by optical microscopy with and without polarizers to reveal birefringence. The images at 50 and 160 °C with and without polarizers are shown in **Figure 6**.

**Figure 6:**
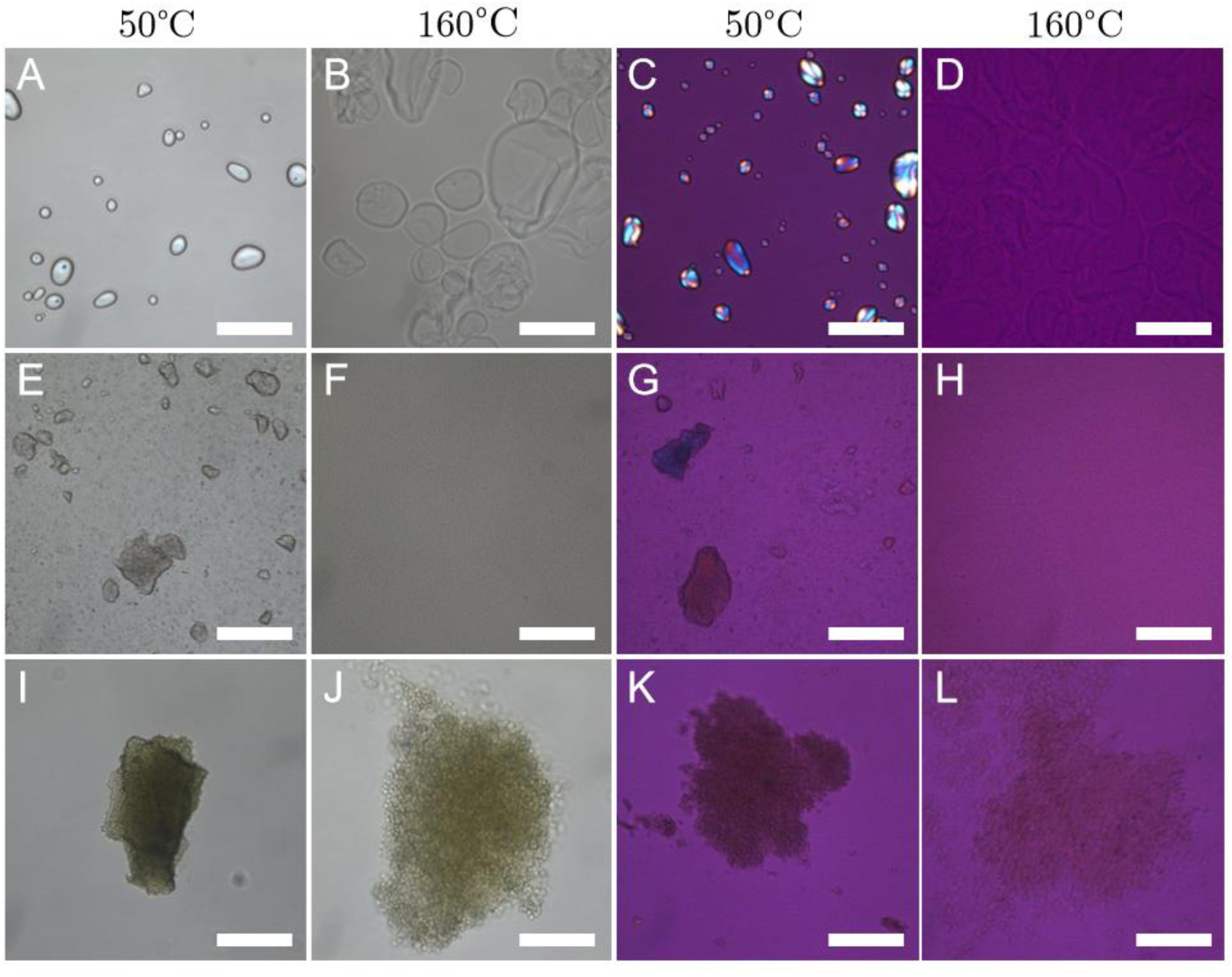
Optical microscopy images of starch granule plasticization in excess glycerol with increasing temperatures. (A-D) Potato starch, (E-H) microalgal starch, and (I-L) starch-rich microalgal cells, observed without (A, B, E, F, I, J) and with polarizers and retardation plate (C, D, G, H, K, L). Scale bar: 100 µm.

The swelling of starch granules with increasing temperature was filmed. In **Video 1** and **Video 2**, potato starch granules considerably swelled at temperatures between 120 and 140 °C, along with a loss of birefringence due to molecular disorganization. At 160 °C, starch granule ghosts were still visible. Conversely, in **Videos 3** and **4**, the swelling of individual microalgal starch granules was hardly visible due to their small size. Nevertheless, agglomerates of starch granules dispersed at around 90 °C, and completely vanished between 120 and 140 °C, forming a homogeneous mixture with no birefringence. On the other hand, **Videos 5** and **6** show that starch-rich microalgal cells remained agglomerated even at elevated temperatures up to 160 °C, but significant swelling by approximately three-fold was observed at both the cell and agglomerate levels. However, the cell walls prevented the observation of the birefringence evolution of starch granules. Therefore, it remained unclear whether the observed cell and agglomerate swelling was attributed to thermal expansion or internal pressure resulting from the swelling and plasticization of starch granules within the cells. In any case, it appears that the actions of the plasticizer and temperature on the whole cells were insufficient to yield a homogeneous, plasticized starch. This supports the need to break down microalgae cell walls beforehand using cell disruption pre-treatments (Günerken, D’Hondt, Eppink, Garcia-Gonzalez, Elst, & Wijffels, 2015), or to break them down directly under the shear forces generated by thermoplastic processing, in order to achieve efficient starch plasticization. In this regard, the starch extraction process implemented in this work (**section 3.1**) appears to be an efficient route.

#### 3.3.2 Plasticization and shaping by extrusion and injection molding

Based on the static plasticization trials, the plasticization ability of the three starch materials was evaluated using a twin-screw extruder. The starch/glycerol batches (70/30 % w/w) were extruded at temperatures of 120 or 140 °C. Interestingly, a green color immediately appeared at the addition of glycerol in the white microalgal starch powder, meaning that microalgal starch was not extensively purified from its chlorophyll. During the extrusion process, microalgal starch did not plasticized properly, leaving a large fraction of the granules in powder form. Since microalgal starch was drier than potato starch, and considering that water is known to aid in starch plasticization, we introduced 10 % water to the microalgal starch before adding glycerol. The final water content for microalgal starch, potato starch and raw biomass was then of 18.4 ± 0.4, 15.7 ± 0.1, and 8.5 ± 0.2 %, respectively. This increased moisture content facilitated proper plasticization of microalgal starch.

The extrudates from the three plasticized starch materials were recovered and subsequently injected into dogbone-shaped molds, as summarized in **Figure S6**. At this stage, a commercial formulation of TPS was introduced as an additional control. Microalgal, potato, and commercial TPS all exhibited proper injection, with important shrinking immediately after unmolding (**Figure 7**). Indeed, the chains of the polymer initially align in the injection direction, but immediately recover their helix configuration when injection is stopped (de Graaf et al., 2003). Interestingly, the shrinkage in microalgal and potato TPS was more pronounced for extrusion temperature of 140 °C than 120 °C. In addition, microalgal TPS displayed a sticky texture after injection and required careful unmolding, unlike the other two TPS. The extrudates produced at 120 and 140 °C from raw microalgae cells were easily injected but did not exhibit the shrinkage behavior typical of TPS.

**Figure 7:**
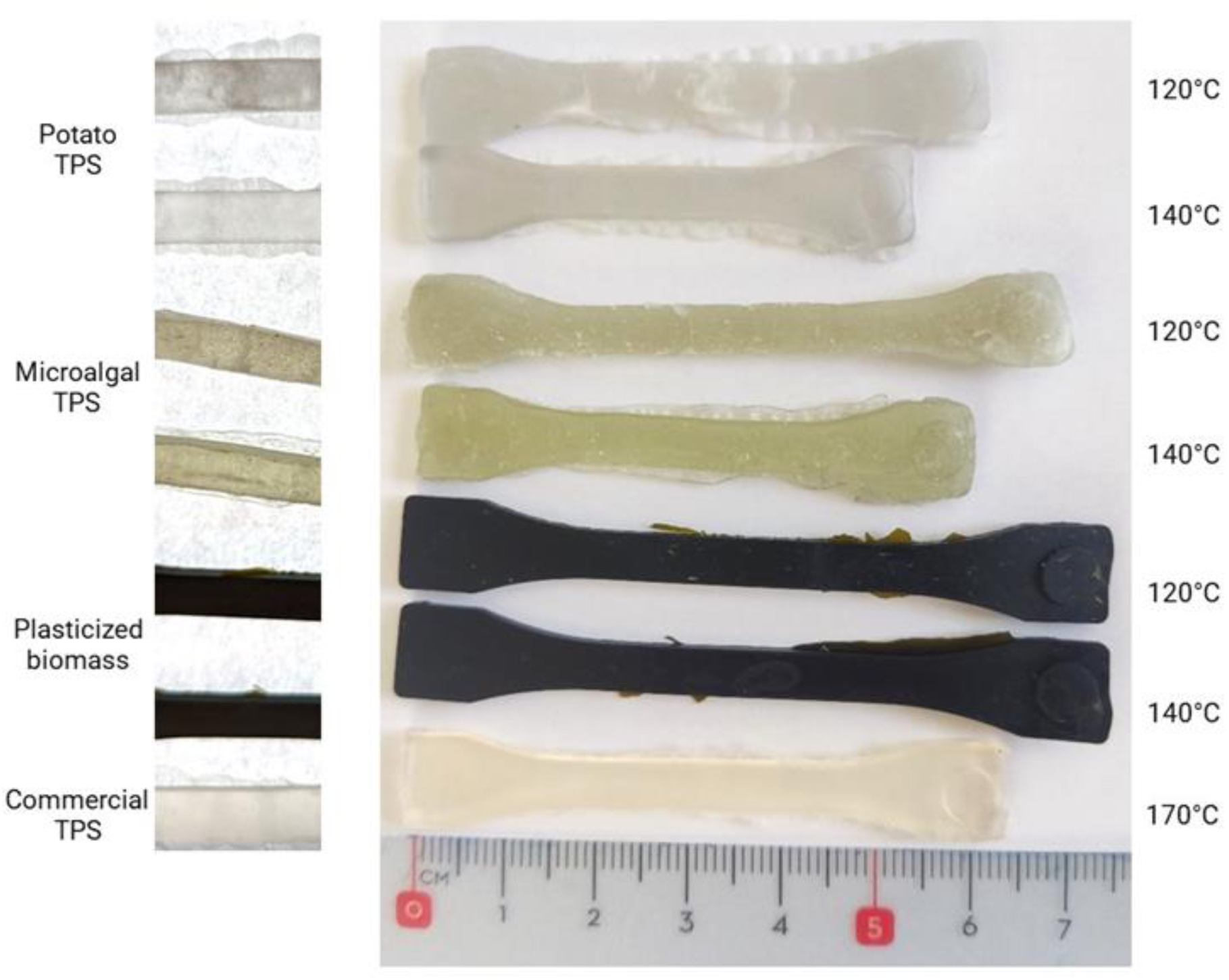
Injected dogbone specimens of potato TPS, microalgal TPS, plasticized raw biomass, and commercial TPS. The dogbone mold measured 7.5 cm in length, but TPS specimens showed immediate shrinkage after unmolding. The temperatures displayed on the right are extrusion temperatures, all injections being performed at the same temperature of 140 °C.

The injected dogbones were cryofractured in liquid nitrogen and were subsequently observed by SEM to study their microstructure (**Figure 8**). Microalgal TPS exhibited proper plasticization from an extrusion temperature of 120 °C, as attested by the homogeneous matrix (**Figure 8B, C**). In contrast, potato TPS displayed good plasticization when extruded at 140 °C, but poor plasticization at 120 °C, characterized by a heterogeneous matrix containing spheroidal particles that likely correspond to ghosts of gelatinized granules (**Figure 8E**). The specimens from entire microalgae cells presented a heterogeneous and granular texture at both extrusion temperatures (**Figure 8H, I**). Closer inspection at higher magnification clearly distinguishes the entire cells, with the starch granules still intact and hence unplasticized (**Figure S7**). In addition, the dogbones of microalgal cells exhibited a significant fragility under traction, highlighting the weakness of intercellular binding. For these reasons, the latter could not be used for tensile tests. In summary, the plasticization of microalgal starch into TPS and its shaping by injection molding was satisfactory, except for the entire microalgae cells and for the potato starch at an extrusion temperature of 120 °C.

**Figure 8:**
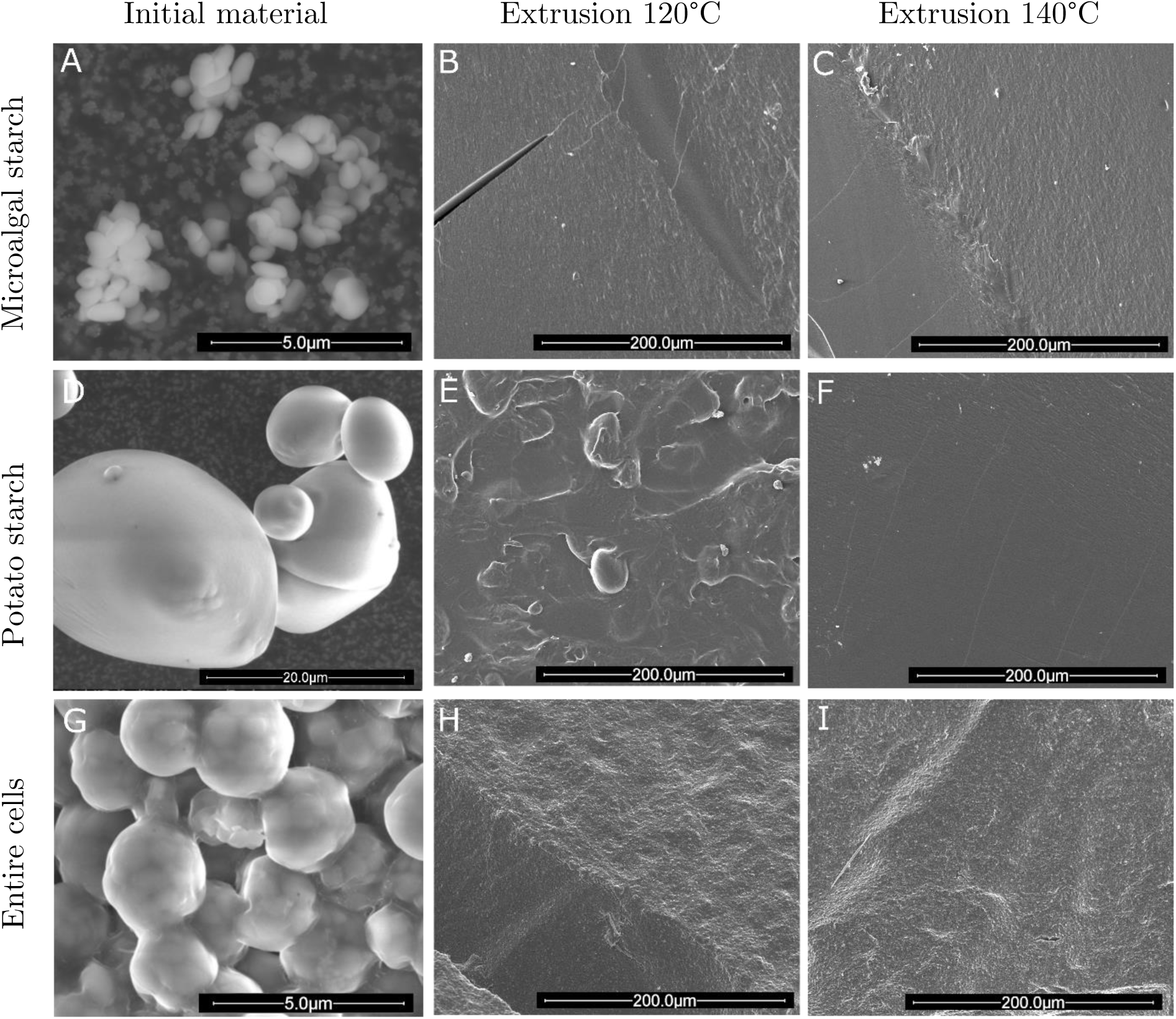
Scanning electron microscopy images of the materials in their initial and plasticized forms. (A-C) Microalgal starch, (D-F) potato starch, and (G-I) entire microalgal cells. The materials are presented in their initial form (A, D, G), extruded at 120 °C (B, E, H), and extruded at 140 °C (C, F, I). (B, C, E, F, H, I) are cross-sections of the cryofractured dogbones. Dogbones were obtained after water addition in algal starch, addition of 30 %w/w glycerol, extrusion, and injection at 140°C.

#### 3.3.3 Tensile mechanical properties

Mechanical properties of injected TPS were studied using tensile tests. In the light of the SEM observations, the extrusion temperature of 140 °C was used to inject additional dogbone specimens of microalgal and potato TPS. After injection and a 3-day resting period, the stiffness, strain at break and strength were analyzed and compared for microalgal, potato, and commercial TPS dogbones. The resulting stress-strain curves are shown in **Figure 9**. First, the reproducibility of the results was very satisfactory with well-grouped stress-strain curves for each material. This supported the fact that the successive processing steps to obtain the TPS specimens, from starch extraction to plasticization by extrusion and injection, have been properly carried out. Secondly, contrasted mechanical behaviors were observed between the different TPS materials. Commercial TPS was more rigid and less ductile, whereas potato and especially microalgal TPS showed a much softer and ductile behavior, with significantly higher elongation but lower strength.

**Figure 9:**
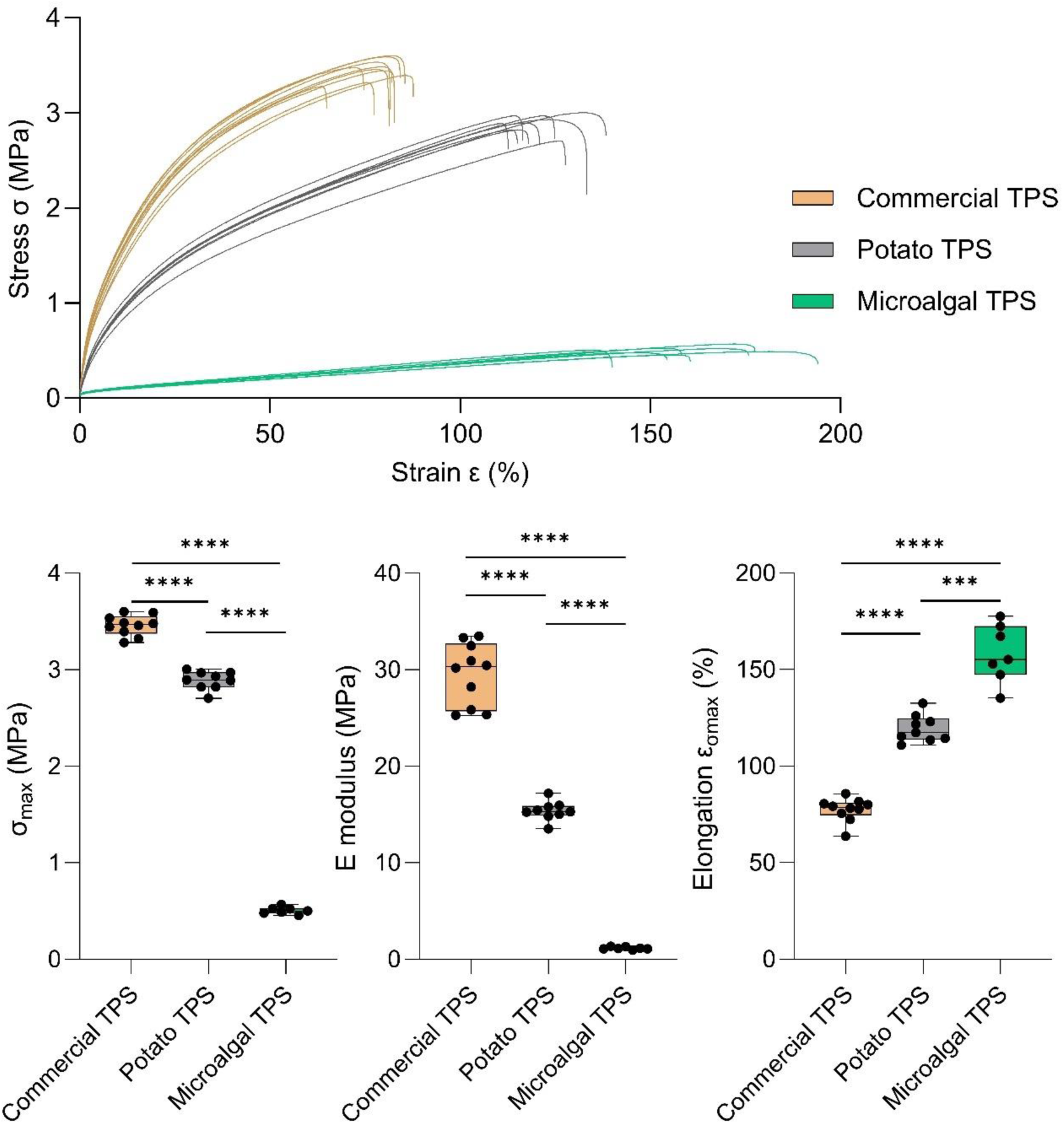
Tensile mechanical properties of commercial, potato, and microalgal TPS evaluated by tensile tests. The p-values for the comparisons indicated in the graph are calculated with Prism 9 (Graphpad Software, LLC) and based on ANOVA Tukey’s multiple comparisons test (***, P = 0.0007; ****, P < 0.0001; 7⩽n⩽10).

The values of maximal stress σ_max_, Young’s modulus E, and elongation ε of potato and commercial TPS were in agreement with the literature (Thunwall, Boldizar, & Rigdahl, 2006; Zhang et al., 2014), and the datasheet provided by the manufacturer, respectively. As pointed out, microalgal TPS was softer and demonstrated much greater elongation compared to potato and commercial TPS, with a mean elongation at maximal stress σ_max_ of 158 % against 120 and 78 %, respectively. Nevertheless, microalgal TPS exhibited maximal stress σ_max_ and E modulus of 0.5 and 1.2 MPa, respectively, which is significantly lower than potato (2.9 and 15.4 MPa, respectively) and commercial TPS (3.5 and 30 MPa, respectively).

A first explanation to the ductile behavior of microalgal starch could be the presence of additional plasticizers. It is known that an increase in plasticizer content in TPS can reduce tensile strength and Young’s modulus while enhancing elongation (Zhang et al., 2014). In our case, the water content was slightly higher in microalgal starch than in potato starch after water addition (18.4 vs. 15.7 %, respectively). Therefore, we reduced the water addition to 7.5 instead of 10 % in microalgal starch (final water content 16.5 %), and produced dogbones for tensile test (**Figure S8**). The mechanical properties were indeed changed, with a slight increase in stiffness and strength, but they did not retrieve the level of potato TPS.

Furthermore, in addition to water and glycerol, various small bound and unbound molecules could act as starch plasticizers, such as mono-hexose (mannose, fructose, glucose), sorbitol, urea or amino-acids (Zhang et al., 2014). Carbohydrates and proteins were present in the initial microalgal biomass **(Table 1**). Possibly, small quantities of small, organic molecules were still present in the extracted starch, contributing to additional plasticizing effect. However, the water rinsing steps during starch purification should have removed any soluble molecules non-bound to starch granules. If present, these plasticizing molecules should be in small amounts, as attested by the final mass balance of purified starch (**Table 1**). Therefore, any plasticizing effect induced by these contaminants shall be limited, as significant changes in mechanical properties typically require to increase the plasticizer content by about 50% (Zhang & Han, 2006).

The second hypothesis explaining the ductile behavior of microalgae TPS involves the potential difference in molecular weight between microalgal starch and potato starch. The molecular weight was reported to significantly influence the mechanical properties of TPS films (Domene-López, García-Quesada, Martin-Gullon, & Montalbán, 2019; Zhang et al., 2014). The average molecular weight of starch depends on its molecular composition. Indeed, the two macromolecular constituents of starch, amylose and amylopectin, exhibit highly different average molecular weights (10^5^ g mol^-1^ and 10^6^-10^7^ g mol^-1^, respectively). For this reason, starches with a high amylose content typically have a lower average molecular weight. Domene-López and coworkers showed that rice, potato, wheat, and corn starches with amylose content of 16.9, 20.5, 24.5, and 24.8 %, respectively, had average molecular weights of 83.2, 69.5, 51, and 51 MDa, respectively (Domene-López et al., 2019).

In the present study, microalgal starch contained 13 % amylose (**Figure 4B**). Standard potato starch was reported to have a typical amylose content of 20 % (Domene-López et al., 2019; Kim, 1996; Malmir, Montero, Rico, Barral, Bouza, & Farrag, 2018). Using the same analytical method as for microalgal starch, the potato starch showed an amylose content of 40 % (two replicates). Therefore, the difference in amylose content is approximately 27 %. As shown in **Figure 9**, potato TPS exhibited higher tensile strength and Young’s modulus, but lower elongation. Domene-López et al. showed that an amylose content higher of 8 % (16.9 % for rice starch vs. 24.8 % for corn starch) resulted in TPS films with approximately two-fold higher tensile strength and Young’s modulus, and three-fold lower elongation (Domene-López et al., 2019). In addition, the impact of amylose content on the mechanical properties of plasticized TPS is linearly valid within the range of 0-30 % amylose (Lourdin, Valle, & Colonna, 1995). Therefore, the difference in amylose content between the microalgal and potato starches, and probably the underlying difference in molecular weight, may have contributed, at least partly, to the observed differences in mechanical properties.

Finally, the impact of crystallinity also needs to be addressed. On the one hand, the difference in crystal type between microalgal and potato starches (A- and B-type, respectively) was not expected to result in a variation in mechanical properties. Indeed, the initial differences in crystalline patterns are lost upon plasticization / extrusion. Domene-López et al. showed that, after plasticization, the crystalline profile of potato TPS was similar to that of rice, wheat and corn TPS, with no discernible impact on mechanical properties (Domene-López et al., 2019). On the other hand, future structural investigations on the amorphous or semicrystalline state of TPS would bring further perspectives on their molecular conformation. In addition, it would be interesting to monitor the evolution of the mechanical properties over time for microalgal TPS. Typically, the amorphous structure of TPS undergoes retrogradation to their crystalline forms during storage (Zhang et al., 2014), especially in humid conditions. This increase in crystallinity might result in significant changes in mechanical properties, resulting in higher tensile strength and Young’s modulus, and lower elongation.

## 4 Conclusion

This study represents a significant step forward in the development of an efficient and industrially relevant starch extraction process from starch-enriched microalgae. This was evidenced by the achievement of high extraction yield and purity degree, reaching 98 and 87 %, respectively. The developed process has also enabled the recovery of valuable fractions such as lipids and proteins, opening up the prospect of a complete biorefinery approach for microalgal biomass.

Microalgal starch granules were successfully plasticized into thermoplastic starch and shaped by injection molding for further analysis of their mechanical behavior. The measured tensile properties were within the range expected for TPS, with much softer, ductile behavior characterized by lower stiffness and strength, but higher elongation.

In this work, we developed a novel, simple and upscalable downstream process for the extraction, purification, and plasticization of microalgal starch. These processes demonstrate the feasibility of recovering microalgal starch with high yields on a pilot scale and indicate that this microalgal starch is suitable for producing TPS. Remarkably, all the developed processing steps can be transposed to an industrial scale. Here, the conversion of raw microalgae into thermoplastic starch on a pilot-scale demonstrates, for the first time beyond the laboratory scale, the relevance of microalgae for the industrial production of bioplastics.

## CRediT

**Alexandre Six:** Conceptualization, Methodology, Validation, Investigation, Writing – original draft, Writing - review & editing. **David Dauvillée:** Investigation, Writing - review & editing. **Christine Lancelon-Pin:** Investigation. **Alexandra Dimitriades-Lemaire**: Investigation, Resources. **Ana Compadre:** Resources. **Carole Dubreuil:** Resources. **Pablo Alvarez:** Project administration, Resources. **Jean-François Sassi:** Project administration, Funding acquisition, Supervision. **Yonghua Li-Beisson:** Supervision, Writing - review & editing. **Jean-Luc Putaux:** Methodology, Investigation, Writing - review & editing. **Nicolas Le Moigne:** Conceptualization, Resources, Writing - review & editing. **Gatien Fleury:** Conceptualization, Methodology, Resources, Funding acquisition, Supervision, Writing - review & editing.

## Acknowledgments

The authors thank Florian Delrue for resources, Sophie Mailley for funding acquisition, Martial Sauceau (IMT Mines Albi) for providing the commercial TPS, Jean-Claude Roux (IMT Mines Alès) for SEM observations, Pierre Sailler and Sonia Molina-Boisseau (CERMAV) for the DSC and granulometry analyses, respectively, as well as the NanoBio-ICMG Platform (UAR 2607, Grenoble) for granting access to the electron microscopy facility. This work was supported by the SEALIVE (https://sealive.eu/) and Nenu2PHAr (https://nenu2phar.eu/) projects, funded by the European Union’s Horizon 2020 Research and Innovation programme under grant agreement #862910 and the Bio Based Industries Joint Undertaking (BBI-JU) under grant agreement #887474, respectively. The BBI-JU receives support from the European Union’s Horizon 2020 Research and Innovation programme and the Bio Based Industries Consortium. The PhD grant of A. Six was partly founded by the French Région Provence-Alpes-Côte-d’Azur (Région Sud).

## Data availability

Data will be made available on request.

## Abbreviations

TPS: Thermoplastic starch

