## Supplementary materials for "From raw microalgae to bioplastics: conversion of *Chlorella vulgaris* starch granules into thermoplastic starch"

### Supplementary material

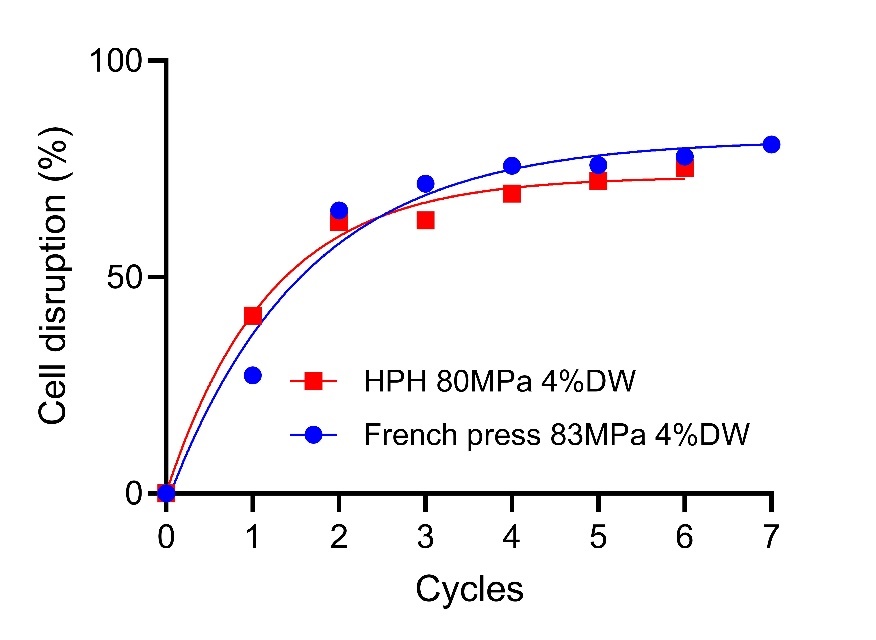

**Figure S1:** Comparison of high-pressure homogenization (HPH) and French press along the pressure cycles for disrupting microalgae cells at similar pressure and concentration. The cell disruption was analyzed in technical triplicates.

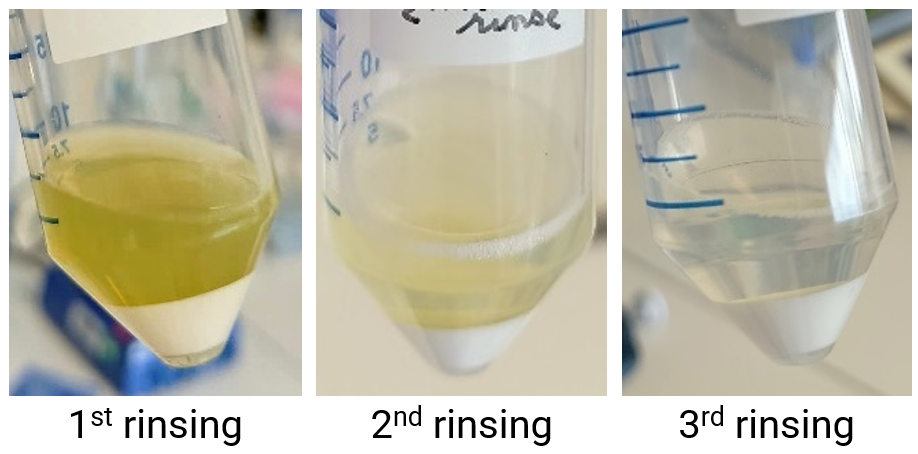

**Figure S2:** Purification of the crude starch pellet consisting in starch resuspension in water, centrifugation, and supernatant removal. The cell debris was eliminated from the supernatant along the successive water rinsing, without using Percoll.

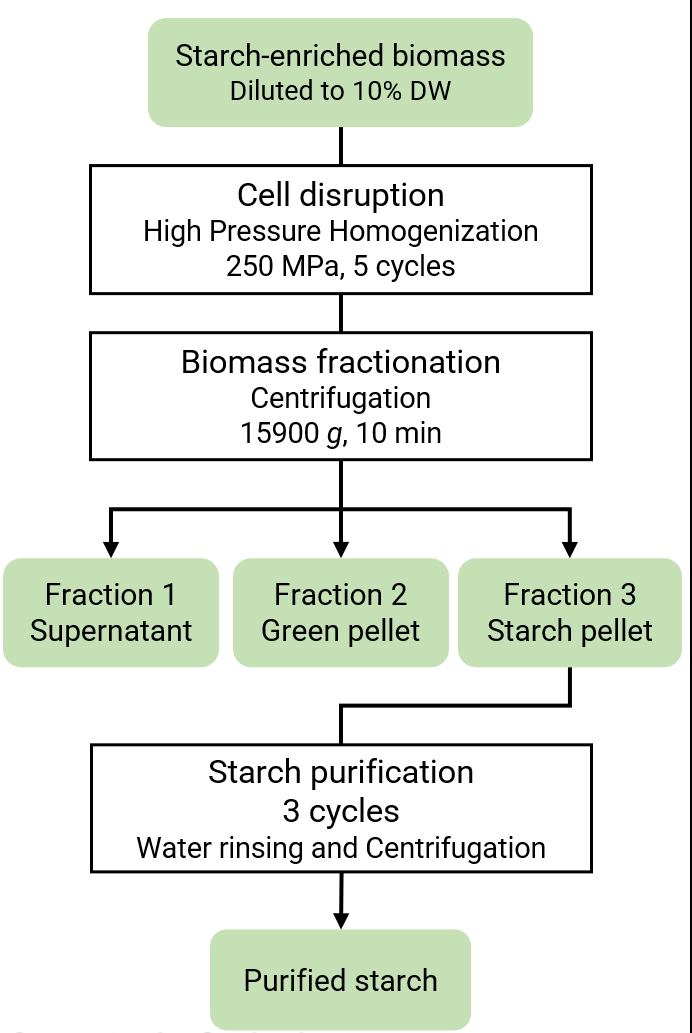

**Figure S3:** Flowchart of the starch extraction protocol.

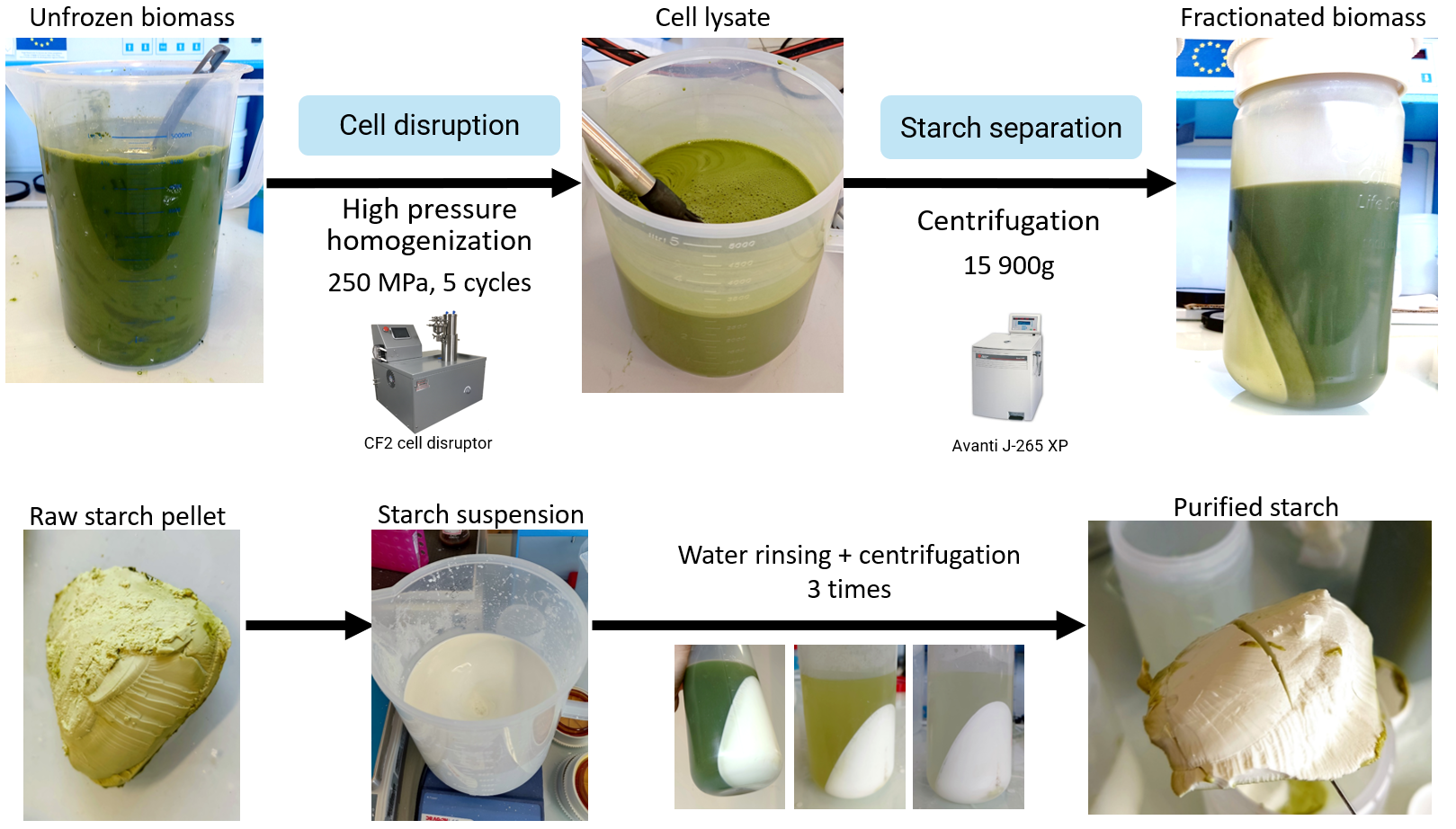

**Figure S4:** Illustration of the starch extraction protocol.

**Table S1:** Recovery of the class of compounds in the different fractions after the separation step. The reported values correspond to the ratio of contents after biomass fractionation (fraction 1, 2, or 3), and before fractionation (cell lysate). The values were normalized to reach a sum of 100 %, as indicated by the factor η.

|  | Dry weight | Starch | Non-glucose carbohydrates | Lipids | Proteins | Ashes |
| --- | --- | --- | --- | --- | --- | --- |
| Fraction 1  Supernatant | 32.3 ± 0.1 % | 3.5 ± 0.5 % | 46.1 ± 9.9 % | 79.5 ± 1.5 % | 61.1 ± 7.9 % | 72.7 ± 4.4 % |
| Fraction 2  Green pellet | 17.2 ± 0.1 % | 2.0 ± 0.6 % | 35.8 ± 7.5 % | 16.3 ± 0.1 % | 36.1 ± 5.2 % | 21.5 ± 1.3 % |
| Fraction 3  Starch pellet | 50.5 ± 1.5 % | 94.5 ± 9.7 % | 18.1 ± 7.0 % | 4.2 ± 0.4 % | 2.7 ± 1.2 % | 5.9 ± 0.5 % |
| Normaliza-  tion factor η | 1.01 | 0.93 | 1.25 | 1.03 | 1.00 | 0.90 |
| Total | 100 % | 100 % | 100 % | 100 % | 100 % | 100 % |

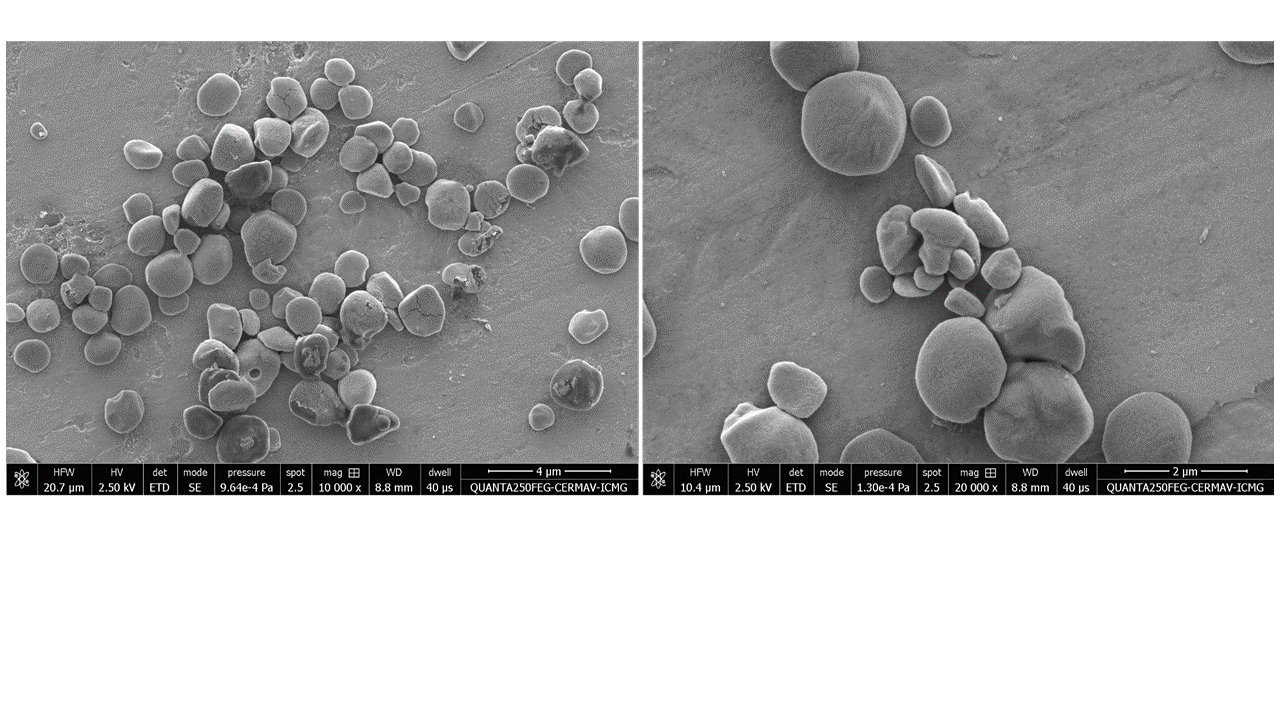

**Figure S5:** SEM images of microalgal starch granules. Their faceted shape was attributed to the tight packing of starch granules in the chloroplast during their accumulation.

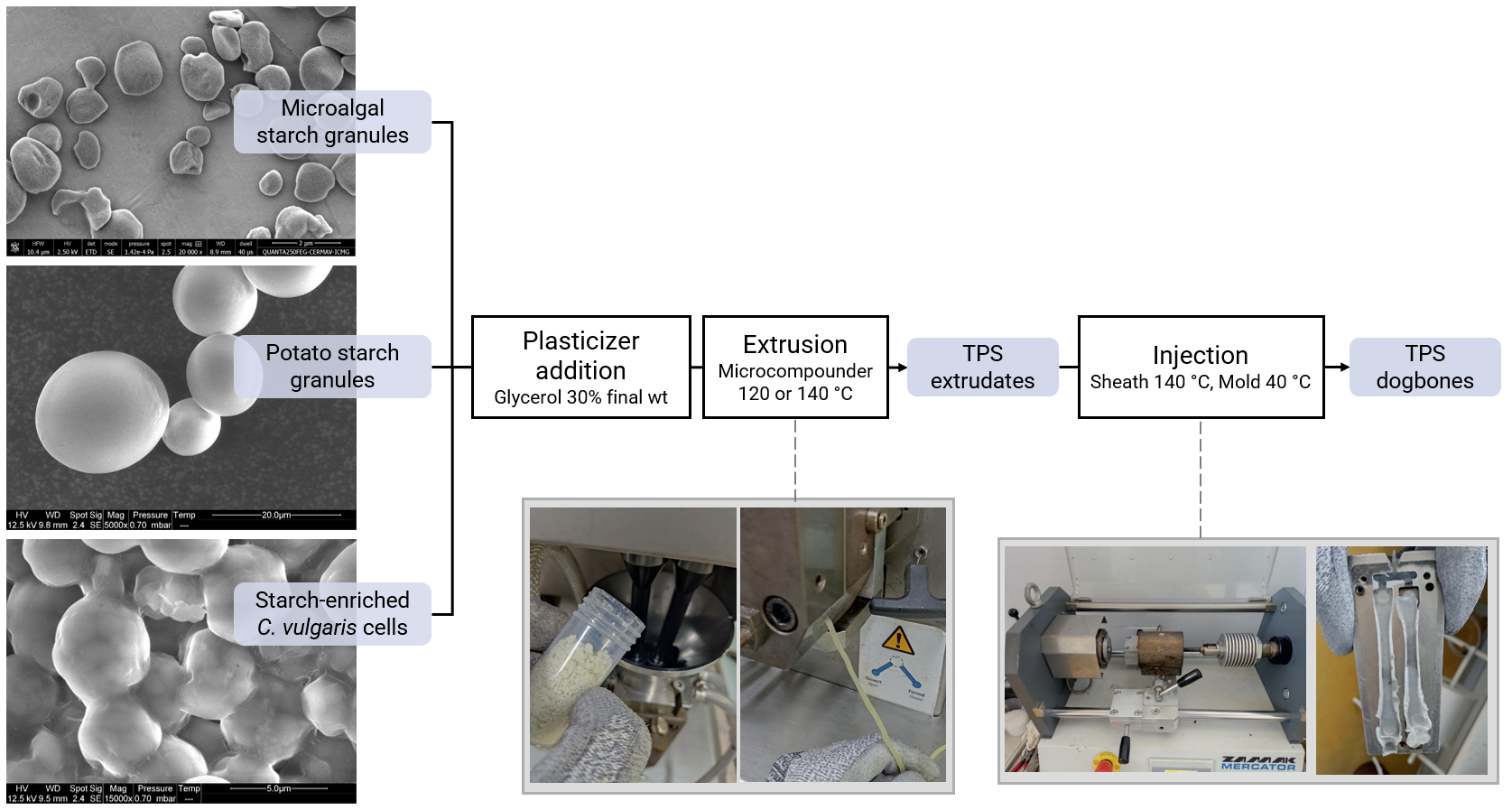

**Figure S6:** Illustration of the plasticization process of starch granules into thermoplastic starch.

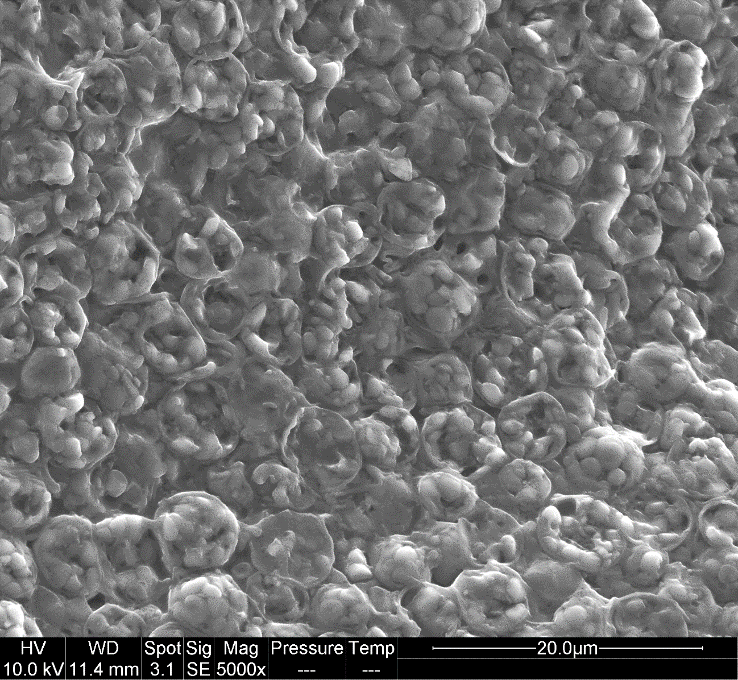

**Figure S7:** Cross-section of a cryofractured dogbone injected specimen of raw biomass. The specimen was prepared by extrusion and injection of a mixture of 70 % w/w entire microalgae cells and 30 % w/w glycerol. Inside the broken microalgae cells, the confined gelatinized starch granules largely retained a spheroidal shape, which indicates that plasticization and homogenization could not be achieved.

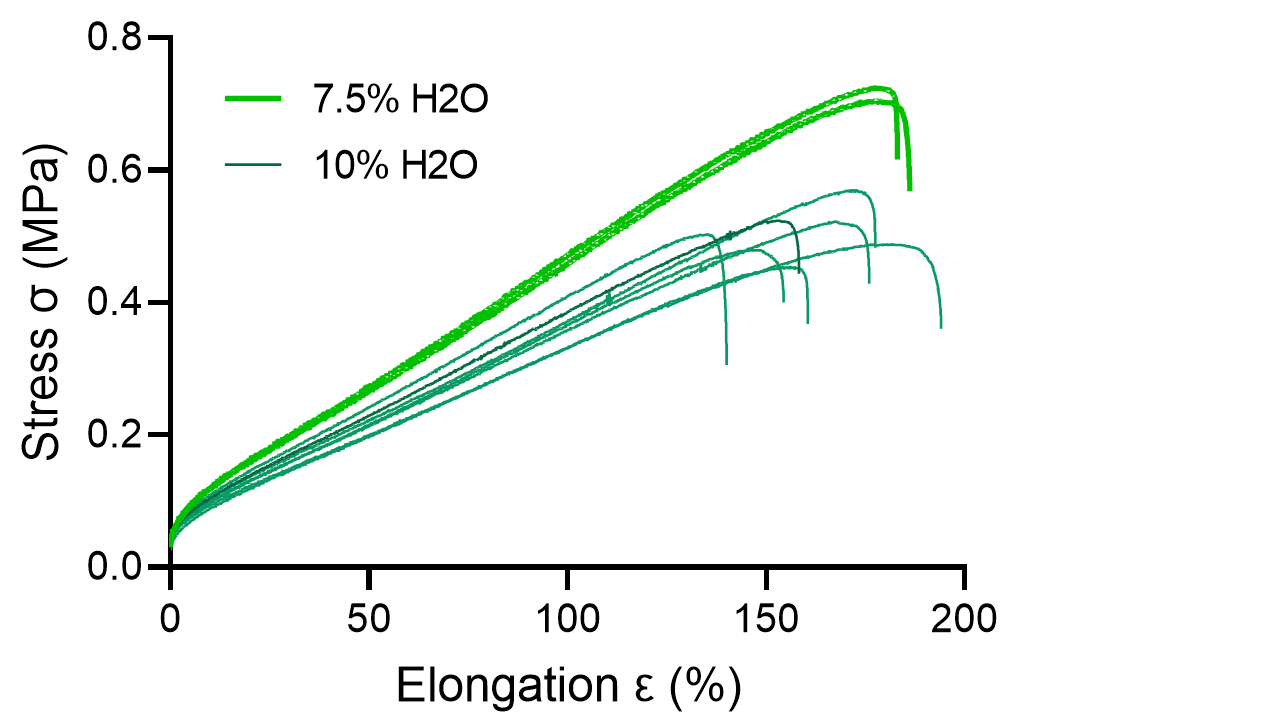

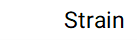

**Figure S8:** Tensile mechanical properties of microalgal TPS with 7.5 or 10 % additional water, evaluated by tensile test.
